# Genomic adaptations to semi-aquatic and aquatic life in spiders

**DOI:** 10.1101/2024.01.15.575295

**Authors:** Zheng Fan, Lu-Yu Wang, Bin Luo, Tian-Yu Ren, Jia-Xin Gao, Piao Liu, Ling-Xin Cheng, Yu-Jun Cai, Bing Tan, Qian Huang, Ming-Qin Deng, Qing Zuo, Xiang-Yun Zhang, Jin-Zhen Lu, Li-Na Sun, Muhammad Irfan, Ning Liu, Chao Tong, Ming Bai, Zhi-Sheng Zhang

**Affiliations:** Key Laboratory of Eco-environments in Three Gorges Reservoir Region (Ministry of Education), School of Life Sciences, Southwest University, Chongqing 400715, China; Key Laboratory of Animal Biodiversity Conservation and Integrated Pest Management, Chinese Academy of Sciences, Beijing 100101, China; Key Laboratory of Zoological Systematics and Evolution, Institute of Zoology, Chinese Academy of Sciences, Beijing 100101, China; University of Chinese Academy of Sciences, Beijing 100101, China; Northeast Asia Biodiversity Research Center, Northeast Forestry University, Harbin 150040, China; School of Life Sciences, Arizona State University, Tempe, AZ 85287, USA

**Keywords:** semi-aquatic and aquatic adaptation, marronoid clade, comparative genomics, molecular evolution, horizontal gene transfer (HGT)

## Abstract

Spiders have primarily adapted to terrestrial life, yet a number of species have made evolutionary transitions to marine and freshwater environments. While its physiological and behavioral adaptations have been characterized, the genetic basis of semi-aquatic and aquatic adaptation in spiders remains poorly understood. Here, we provide a high-quality, chromosome-level genome assembly for the aquatic spider *Argyroneta aquatica*, alongside a reference transcriptome for the semi-aquatic spider *Desis martensi*. We performed comparative genomes analyses of 22 spider species, including a unique aquatic spider, two semi-aquatic spiders and 19 terrestrial spider species, with a focus on those in the marronoid clade. By integrating morphological, phylogenomic, comparative genomic, transcriptomic, and metabolomic analyses, we explored the genomic adaptations of aquatic and semi-aquatic spiders. Phylogenomic analysis suggests that aquatic and semi-aquatic spiders have independently evolved from their terrestrial ancestors and represent divergent evolutionary routes We found hundreds of genes tend to experience relaxed selection, positive selection, and evidence of horizontal gene transfer (HGT) associated with the transition to aquatic and semi-aquatic life in spiders. These genes are associated with respiratory, osmoregulatory, fat metabolism and digestion, hypoxia, and thermal functions, putatively facilitate the adaptations to diverse underwater life. Altogether, our findings highlight the divergent evolutionary mechanisms enabling spiders to thrive in diverse aquatic environments, providing insights into the genomic basis of adaptations to semi-aquatic and aquatic habitats.

## Introduction

Most spiders have adapted to terrestrial life, while a noteworthy proportion of spider species, which were originally terrestrial, have undergone a reversal of adaptation, back to marine and freshwater habitats (Leggett, Vink, and Nelson, 2024). This evolutionary transition has occurred either in semi-aquatic or fully aquatic spider species. One exemplar group is the marronoid clade, encompassing approximately nine families including Agelenidae, Amaurobiidae, Cybaeidae, Cycloctenidae, Desidae, Dictynidae, Hahniidae, Stiphidiidae, and Toxopidae, which exhibit diverse life-history strategies, particularly adapted to a range of extreme environments (Gorneau et al., 2023). For instance, spiders that traditionally inhabited terrestrial environments have demonstrated semi-aquatic lifestyle, such as those in the intertidal spider genus *Desis* (Desidae), which are found on rocky coasts worldwide. Notably, the unique water spider *Argyroneta aquatica* (Dictynidae) has undergone complete adaptation to an aquatic lifestyle (Stefano et al., 2016; Schaber et al., 2023).

Various physiological adaptations to semi-aquatic and aquatic environments have been revealed in spiders. Respiration is most challenging for semi-aquatic and aquatic spiders, because their respiratory systems are the same as these in terrestrial spiders, with book lungs and trachea. The semi-aquatic spider *Desis* live in the spaces among the rocks and crevices and/or shells with available air space in and around the nest (McQueen et al., 2012). While the water spider can construction a “diving bell” for breath underwater, which is filled with air transported from the water surface (Schaber et al., 2023). Due to limited air availability, hypoxic adaptation is especially important for aquatic spider (Seymour and Hetz, 2011). For semi-aquatic spiders *Desis* living in marine habitats, osmoregulation represents one of the most important physiological processes. In semi-aquatic spiders, a variety of physiological, behavioral, or morphological adaptations are caused by osmotic pressure. Unlike terrestrial spiders which prey on insects, fish may represent one of the important food sources for semi-aquatic and aquatic spiders (Nyffeler and Pusey, 2014). Thus, many morphological, physiological, and behavioral characteristics differ among terrestrial, semi-aquatic, and aquatic spiders. However, the molecular and genomic differences underlying adaptations to semi-aquatic and aquatic life in spiders remain poorly understood due to the long-term lack of relevant genomic resources.

The divergent adaptations of spiders to different habitats are hypothesized to be associated with several evolutionary mechanisms, including relaxed selection, positive selection, and likely horizontal gene transfer (HGT). Specifically, species tended to undergo diverse selections on key phenotypic traits during the adaptation to a new environment. Once these key phenotypic traits reach a new optimal value, they may tend to experience weaker selection, leading to a relaxation of selective pressures (Hughes, 2012; Yoder et al., 2010; Hunt et al., 2011). For instance, cetaceans (whales and dolphins) exhibit dramatic morphological changes from their terrestrial ancestors, including the development of flippers and tail flukes, driven by rapid evolutionary processes (Yim et al., 2014; Tian et al., 2016; Noh et al., 2022). In addition, positive selection can drive the fixation of beneficial mutations that enhance fitness in a particular habitat, as seen in a cold-adapted ant (*Tetramorium alpestre*) (Cicconardi et al., 2020). Another example, bdelloid rotifers are microscopic aquatic animals that have acquired a significant portion of their genome through HGT from bacteria, fungi, and plants, which involved in various metabolic processes, including the degradation of complex carbohydrates and the synthesis of essential amino acids, which facilitate the rotifers surviving in nutrient-poor aquatic environments (Gladyshev et al., 2008). Increasingly, HGT has been documented in insects, where it can contribute to the acquisition of genes that offer survival advantages in specific environments (Neumann et al., 2013). In *Pyrrhocoris apterus* (Pyrrhocoroidea), salivary proteins putatively derived from HGT are critical for salivary sheath formation and other feeding processes (Huang et al., 2024). The horizontally acquired pectin-digesting polygalacturonases (PGs) of the leaf beetle *Phaedon cochleariae* from fungi, which enhance nutrient accessibility, development, and survival (Kirsch et al., 2022). In silkworms and ladybird beetles, HGT-acquired genes have been identified that contribute to pathogen resistance (Zhu et al., 2011; Li et al., 2021). In lepidopteran insects, evidence suggested that HGT-acquired genes is linked to male courtship behavior (Li et al., 2022). Moreover, in whitefly and leaf beetle, HGT-acquiredgenes are essential for survival (Gilbert and Maumus, 2022). The SMaseD enzyme, a distinctive toxin enzyme isolated from the spider *Loxosceles*, has been suggested to evolve through HGT, providing the initial evidence of this phenomenon in spiders (Binford et al., 2005). However, little is known about whether these evolutionary mechanisms contribute to the divergent adaptation to semi-aquatic and fully aquatic life in spiders.

In this study, we generated a high-quality chromosome-scale genome assembly for the aquatic spider *Argyroneta aquatica* and a reference transcriptome for the semi-aquatic spider *Desis martensi*, and integrated publicly available genomes and transcriptomes of their closely related spider species. We performed comparative genomics and molecular evolution analyses of above 22 spider species, including the unique aquatic spider, two semi-aquatic spiders and 19 terrestrial spider species within the in the marronoid clade. Moreover, we presented the use of comparative transcriptomics and metabolomics with the focus on hypoxia response in water spiders. Altogether, we sought to identify genomic signatures associated with semi-aquatic and aquatic adaptations in spiders.

## Results

### Spider genomic features and phylogenomic analyses

To investigate the genomic adaptations to semi-aquatic and aquatic life in spiders, we selected 22 species, including the unique aquatic spider species, *Argyroneta aquatica*, two semi-aquatic spider species (*Desis martensi* and *Desis japonica*), and 19 terrestrial spider species (*Uloborus diversus* as outgroup) (Fig. 1A and 1B, Table S1) for comparative analyses, with the focus on those in the marronoid clade. In this study, we sequenced the genome of water spider *A. aquatica* and a reference transcriptome of one semi-aquatic species *D. martensi*. We also re-annotated the publicly available genomes of two closely related terrestrial spider species that does not have RefSeq annotations, and *de novo* assembled 17 species based on raw RNA sequencing data from NCBI database (Table S1). Specifically, we reported a high-quality and chromosome-level genome of water spider *A. aquatica*, with an N50 of 2.44 Mbp and total assembly length of 753.77 Mbp that distributed in 11 chromosomes (Fig. 1C, Fig. S1, Table S2), detecting 98.0% arthropod conserved genes (1,013 genes) by BUSCO (Table S2). In addition, we found that approximately 41.53% of the whole genome contains repetitive regions (Fig. 1C, Table S3). A combination of homolog search, transcriptome-based and *ab initio* prediction for searching gene models resulted in a total of 17,632 protein-coding genes (Table S4). Moreover, building on the existing transcriptome of a semi-aquatic spider, *D. japonica*, we supplemented a reference transcriptome of another semi-aquatic species, *D. martensi*, detecting 97.2% arthropod conserved genes by BUSCO. We performed genome re-annotation for two closely related terrestrial spider species, *Amaurobius ferox* (GCA_951213105.1) and *Dolomedes plantarius* (GCA_907164885.2), resulting in 14,258 proteins for *A. ferox* with BUSCO completeness scores of 74% and 76,921 proteins for *D. plantarius* with 80.6% arthropod conserved genes (Table S4).

**Figure 1.**
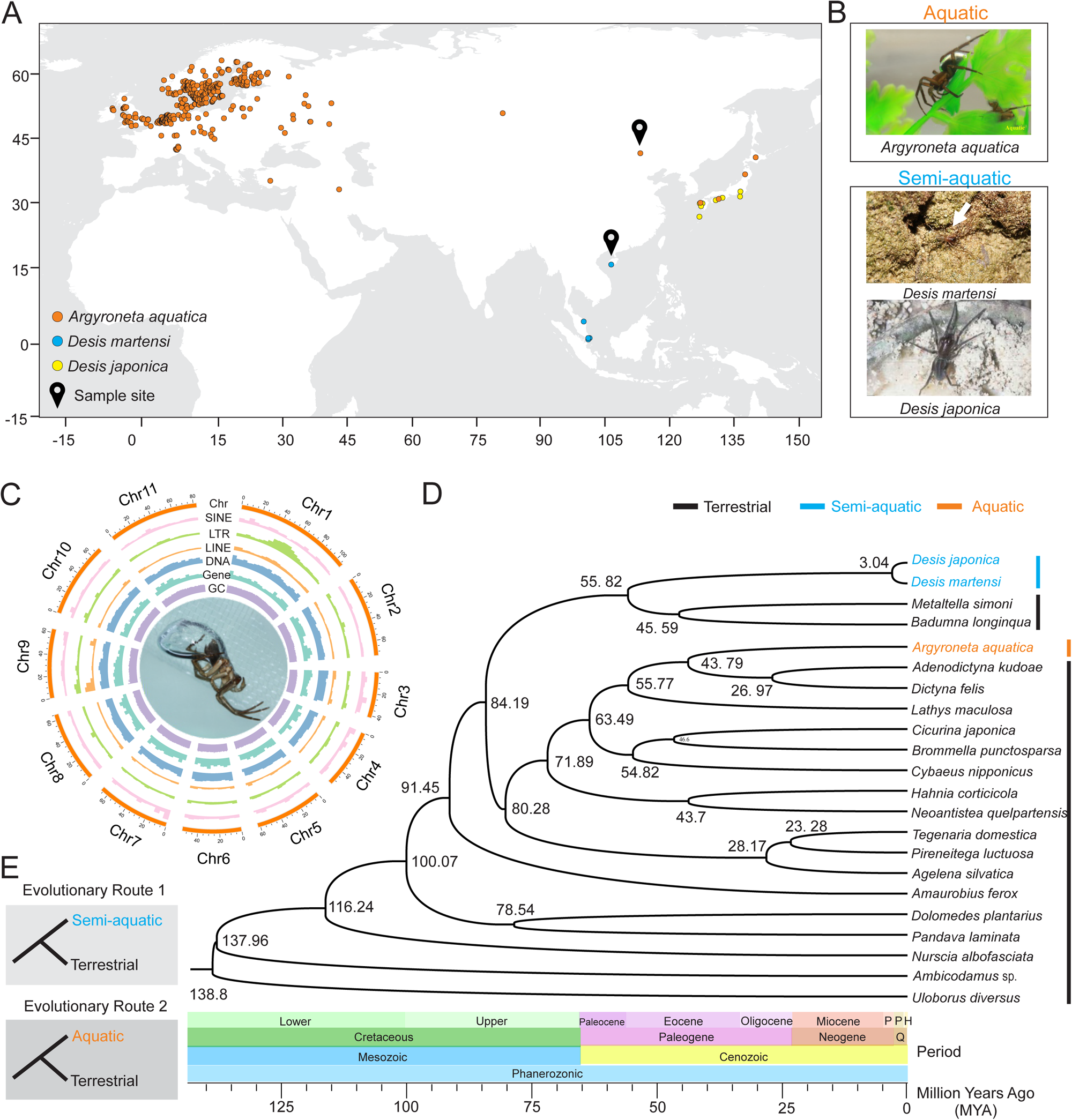
The genome structure analysis and phylogenetic analysis of water spider *Argyroneta aquatica* with other spiders. A. The global distribution of *A. aquatica* and *Desis martensi* and *Desis japonic*. The distribution data for aquatic (*A. aquatica*) and semi-aquatic (*D. martensi* and *D. japonica*) spiders from the World Spider Catalog (https://wsc.nmbe.ch/). Based on the latitude and longitude coordinates information, the point map for global distribution of aquatic and semi-aquatic spiders drawing by the online website of SimpleMappr (https://www.simplemappr.net/#tabs=0). B. Images of *A. aquatica, D. martensi* and *D. japonic.* The Photograph of *A. aquatica* by Qian-Le Lu, *D. martensi by* Lu-Yu Wang. And the Photograph of *D. japonic* was cited from Ono (Ono, 2009). C. Distribution of chromosomal elements of water spider. The inner ring contains a picture of water spider. The outer rings of the circle represent means bellow, respectively: Chr: chromosomes, SINE: short interspersed nuclear element, LTR: long terminal repeat, LINE: long interspersed nuclear elements, DNA: DNA transposable elements, Gene: distribution of genes, GC: GC content. The Photograph of *A. aquatica* by Kun Yu. D. Phylogenetic analysis of *A. aquatica* and most “marronoid” spider species with the Uloboridae (*U. diversus*) as outgroups. The divergence times among different species are shown at the bottom and branch node. The black line represents the spiders’ terrestrial living habits, blue line with semi-aquatic living habits, and orange line with aquatic living habits. E. The hypothesis of evolutionary route of terrestrial, Semi-aquatic and aquatic.

The genome-scale phylogeny of aquatic spider, *A. aquatica*, semi-aquatic spiders *D. japonica*, *D. martensi*, and other 19 terrestrial spiders were reconstructed based on a set of 941 single copy orthologous genes. Molecular dating analysis indicated that the divergence time of marronoid spider clade was split approximately 100.07 million years ago (Fig. 1D). The aquatic spider *A. aquatica* is a sister group to Dictynid spiders (i.e. *Adenodictyna kudoae*, *Dictyna felis* and *Lathys maculosa*) with the divergence time at 43.79 million years ago. The *Desis* (*D. japonica* and *D. martensi*) as members of the family Desidae with semi-aquatic lifestyle, while species within the other genera of this family (i.e. *Metaltella simoni* and *Badumna longinqua*) with terrestrial lifestyle, with the divergence time at 55.82 million years ago (Fig. 1D).

### Proposed Semi-aquatic and aquatic transitions

We proposed the hypotheses of the divergent evolutionary routes for semi-aquatic and aquatic spiders (Fig. 1E). Semi-aquatic and aquatic spiders have each independent route evolved from ancestral terrestrial spiders. First, water spiders inhabit freshwater lakes or ponds, while semi-aquatic spiders (*M. simoni* and *B. longinqua*) live in intertidal zones along the seashore (Fig. 1A, 1B). Phylogenetic analysis revealed that semi-aquatic spiders, along with closely related species (e.g. *M. simoni* and *B. longinqua* ) within the same family, exhibit a terrestrial lifestyle. Similarly, aquatic spider, along with its closely related species (e.g. *A. kudoae*, *D. felis*, and *L. maculosa*) within the same family, also exhibit a terrestrial lifestyle (Fig. 2A, 2B).

**Figure 2.**
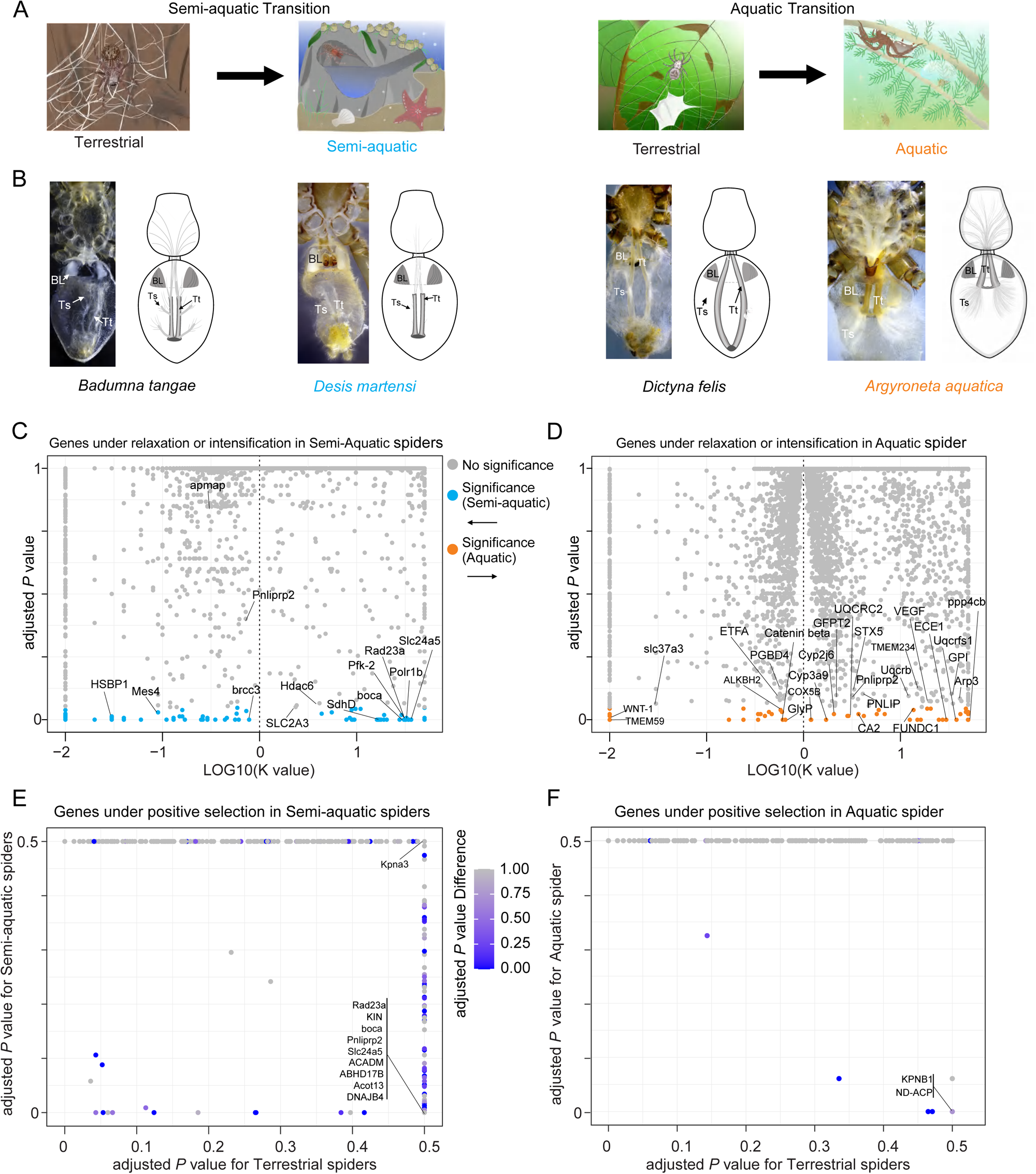
The environment, respiratory systems, and genome-wide dN/dS changes of spiders transitioning from terrestrial to semi-aquatic and terrestrial to aquatic environments. A. Diagram illustrating the transition of spiders from terrestrial to semi-aquatic and terrestrial to aquatic environments. B. The morphological of respiratory systems changes of spiders transitioning from terrestrial to semi-aquatic and terrestrial to aquatic environments. Tt: tracheal tube; Ts: tracheoles; BL: book lungs. Tracheoles are extremely fine tubes, often less than 1 micron in diameter. C. The volcano plot of genes under relaxation or intensification selection of semi-aquatic spiders compared with terrestrial spiders. D. The volcano plot of genes under relaxation or intensification selection of aquatic spiders compared with terrestrial spiders. E. The volcano plot of genes under positive selection of semi-aquatic spiders compared with terrestrial spiders. F. The volcano plot of genes under positive selection of aquatic spiders compared with terrestrial spiders.

### Relaxed selection associated with spider underwater transitions

We estimated the gene-wide ratios of non-synonymous to synonymous substitution (dN/dS, ω) across the spider phylogeny for 4,744 (semi-aquatic transition) and 4,871 (aquatic transition) of shared orthologs, and detected the selection acting on two divergent underwater transitions in spider by calculating three discrete categories of ω ratios (purifying selection: ω1, neutral evolution: ω2, positive selection: ω3) using RELAX. The parameter K-value indicated overall relaxation (K < 1) or intensification (K > 1) of selection.

For semi-aquatic transition, we identified genes that experienced relaxed or intensified selection in semi-aquatic species relative to the other species involved in “evolutionary route 1” (Fig. 1E). We found 58 genes under intensification of selection (K > 1, q-value <0.05), and 67 genes under relaxation of selection (K < 1, q-value <0.05) (Table S5). The sets of genes under intensification and relaxation were not associated with statistically significant enrichment of specific biological functions (Fig. S2-S5). In semi-aquatic spiders, among these genes under intensified selection, we found a set of genes associated with respiratory chain and energy metabolism, such as *SdhD* (Succinate dehydrogenase [ubiquinone] cytochrome b small subunit), *Pfk-2* (phosphofructokinase-2/fructose-2,6-bisphosphatase), *SLC2A3* (Solute Carrier Family 2, Facilitated Glucose Transporter Member 3), with trachea devolopment, such as *Hdac6* (Histone deacetylase 6), with transmembrane ion transport, such as *Slc24a5* (Sodium/potassium/calcium exchanger 5), with fat metabolism and digestion, such as *boca* (LDLR chaperone boca), *Pnliprp2* (Pancreatic lipase-related protein 2), with DNA repair such as *Polr1b* (DNA-directed RNA polymerase I subunit RPA2) and *Rad23a* (UV excision repair protein RAD23 homolog A) (Fig. 2C). Among these genes under relaxation of selection, we found genes associated with fat metabolism and digestion, such as *apmap* (Adipocyte plasma membrane-associated protein), with respond to heat stress such as *HSBP1* (Heat shock factor-binding protein 1), with DNA repair such as *brcc3* (Lys-63-specific deubiquitinase BRCC36) and *Mes4* (DNA polymerase epsilon subunit 4) (Fig. 2C).

For aquatic transition, we identified genes that experienced relaxed or intensified selection in water spider species relative to the other species involved in “evolutionary route 2” (Fig. 1E). We identified 92 genes under intensification of selection (K > 1, q-value <0.05), and 59 genes under relaxation of selection (K < 1, q-value <0.05) (Table S6). The sets of intensification genes were associated with statistically significant enrichment of nucleolus, obsolete chromosomal part, and small molecule catabolic process (Fig. S6, S7). Among the set of genes under intensification of selection, we found genes may be associated with the mitochondrial electron transport chain, such as *Uqcrfs1* (Cytochrome b-c1 complex subunit Rieske), *COX5B* (Cytochrome c oxidase subunit 5B), *UQCRC2* (Cytochrome b-c1 complex subunit 2), *FUNDC1* (FUN14 domain-containing protein 1), *CA2* (Carbonic anhydrase 2), and *Uqcrb* (Cytochrome b-c1 complex subunit 7), with the development of the tracheae such as *TMEM234* (Transmembrane protein 234 homolog), *Arp3* (Actin-related protein 3) and *ECE1* (Endothelin-converting enzyme 1), *STX5* (Syntaxin-5), *VEGF* (Vascular endothelial growth factor A), with energy metabolism such as *GPI* (Glucose-6-phosphate isomerase), *GFPT2* (Glutamine--fructose-6-phosphate aminotransferase) and *UQCRC2* (Cytochrome b-c1 complex subunit 2, mitochondrial), with cytochrome P450 enzymes such as *Cyp2j6* (Cytochrome P450 2J6) and *Cyp3a9* (Cytochrome P450 3A9), with DNA repair such as *ppp4cb* (Serine/threonine-protein phosphatase 4 catalytic subunit B), with fat metabolism and digestion such as *Pnliprp2* (Pancreatic lipase-related protein 2) and *PNLIP* (Pancreatic triacylglycerol lipase). While the sets of genes under relaxation were not associated with statistically significant enrichment of specific biological functions (Fig. S8, S9). Among the set of genes under relaxation of selection, we found genes may associated with the development of lung and trachea such as *Wnt-1, Catenin beta, TMEM59* (Transmembrane protein 59), with energy metabolism such as *GlyP* (Glycogen phosphorylase), *slc37a3* (Sugar phosphate exchanger 3), *ETFA* (Electron transfer flavoprotein subunit alpha, mitochondrial ), with DNA repair such as *ALKBH2* (DNA oxidative demethylase ALKBH2), *PGBD4* (PiggyBac transposable element-derived protein 4) (Fig. 2D).

### Positive selection facilitates spider adaptations to underwater life

We identified 127 positively selected genes (PSGs) in semi-aquatic spiders (Table S7). Among these genes, we found some genes associated with transmembrane ion transport such as *Slc24a5* and *slc37a3,* with DNA repair such as *KIN* (DNA/RNA-binding protein KIN17) and *Rad23a* (UV excision repair protein RAD23 homolog A), with fat metabolism and digestion such as *Pnliprp2, boca, ACADM* (Medium-chain specific acyl-CoA dehydrogenase, mitochondrial), *ABHD17B* (Alpha/beta hydrolase domain-containing protein 17B)*, Acot13* (Acyl-coenzyme A thioesterase 13), with respond to heat stress such as *DNAJB4* (DnaJ homolog subfamily B member 4), with respiratory chain and energy metabolism such as *SdhD* (Succinate dehydrogenase), *SLC2A3*, with DNA repair such as *KIN* (DNA/RNA-binding protein KIN17), with karyopherin gene *Kpna3* (Importin subunit alpha-4) (Fig. 2E). While, a total of 26 PSGs were identified in aquatic spider (Table S8), which were associated with fat metabolism and digestion such as *ND-ACP* (*Acyl carrier protein, mitochondrial*), or with karyopherin gene *Kpnb1* (Importin subunit beta-1) (Fig. 2F).

### Divergent morphology of respiratory systems

The book lung and trachea of four spider species were dissected, respectively. Significant morphological differences between semi-aquatic spider (*D. martensi*) and its terrestrial relative (*Badumna tangae*) within semi-aquatic transition, aquatic spider species (*A. aquatica*) and its terrestrial relative (*D. felis*) within aquatic transition had been observed in their book lung (Fig. 2B). Specifically, compared to its closely related terrestrial species *B. tangae*, the semi-aquatic *D. martensi* has more tracheoles and thicker book lungs. The respiratory system of the water spider (*A. aquatica*) shows significant variation. Compared to its terrestrial relative *D. felis*, water spider has spiracles located closer to the front of its body, along with more tracheoles and thicker book lungs (Fig. 2B).

### Expansion of the ABC and ACAD gene families through putative horizontal gene transfer (HGT)

Horizontal gene transfer (HGT) has been hypothesized as one of evolutionary mechanisms may facilitate the underwater adaptation in water spider. The ABC and ACAD gene families in water spider genome were found to be significantly expanded in comparison to terrestrial spider *A. ferox* and *D. plantarius* genomes (Fig. 3A, 3B). In addition, a set of genes belong to ABC and ACAD gene families that are unique to water spiders showed highly homologous with bacteria. We further validated the reliability of inference of these putative HGT-acquired genes (i.e. ABC and ACAD gene families) using a combination of approaches including Blast-search, phylogenetic mapping, accessing RNA expression, whole genome resequencing reads mapping, and PCR cloning (see methods for details). In total, we identified 75 ABC genes in water spider genome, including 39 genes acquired through HGT. While, its closely related terrestrial species *A. ferox* has 26 ABC genes, and *D. plantarius* has 41 ABC genes (Table S9). For ACAD genes, we identified 35 ACAD genes in water spider genome, including 21 genes acquired through HGT. While its closely related species *A. ferox* has 9 ACAD genes, and *D. plantarius* has 12 ACAD genes (Table S10). To further understand the characteristics of HGT-acquired genes in ABC and ACAD gene families of water spider, we analyzed the number of Exons and the GC content of each gene family member (Fig 3A, 3B, 3C, Table S9, S10). Intriguingly, we found that the HGT genes have only 1-2 exons, which is significantly fewer than the non-HGT genes. The GC content of the HGT genes varies widely, differing from that of the non-HGT genes. This variation may be because the GC content of the HGT genes is related to the GC content of the source bacteria. The distribution of HGT genes was on scaffolds that have not been assembled into chromosomes. Additionally, the distribution of these HGT genes is relatively scattered (Fig. 3D and 3E). We mapped the HGT genes of ABC and ACAD gene families to the genome resequencing data of water spider. Except for ABCF-15 (gene ID number: Argyroneta_aquatica_00016198), most genes can be mapped to the genome re-sequencing reads (Fig. 3F).

**Figure 3.**
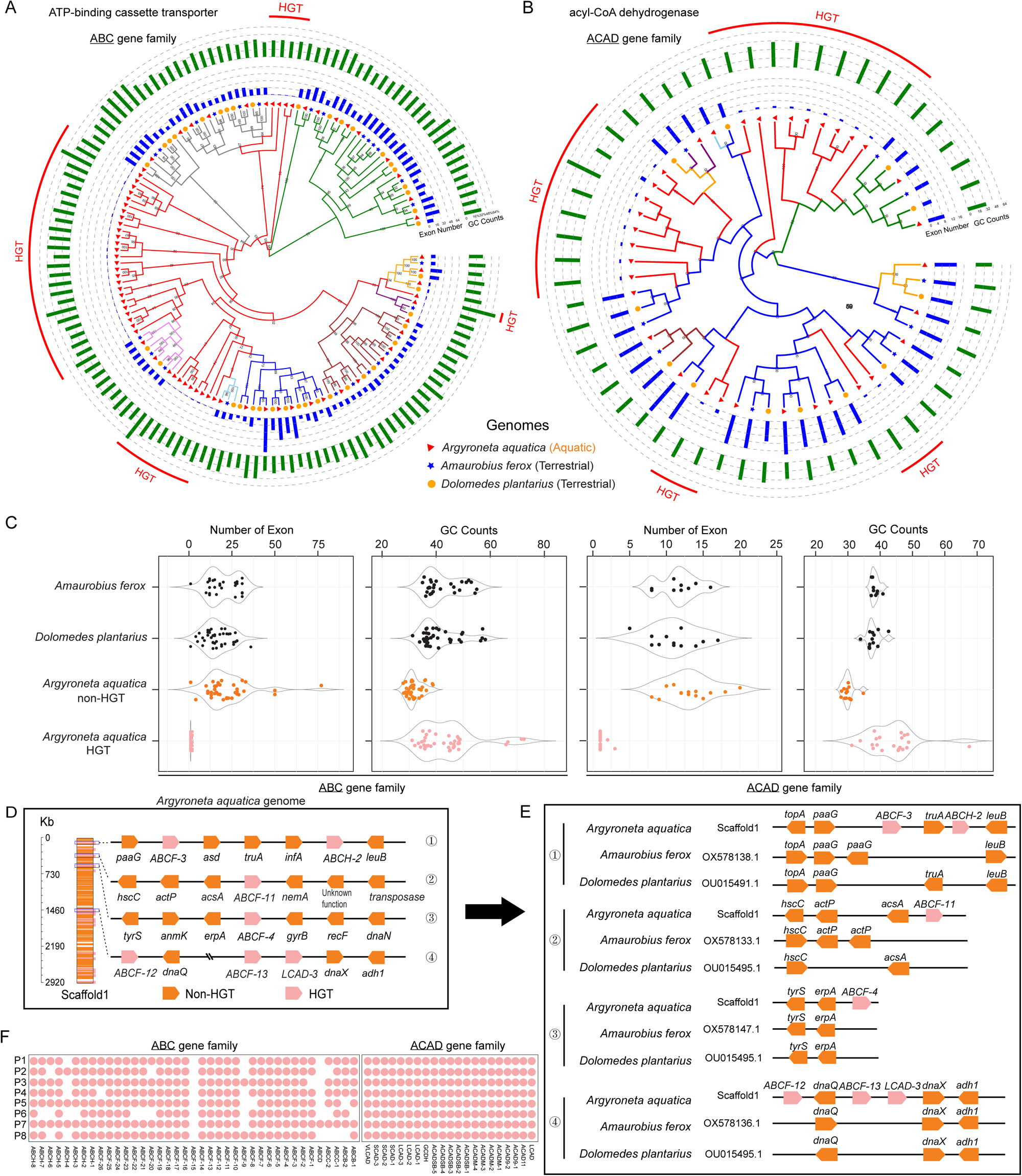
The feature of HGT-acquired ABC and ACAD genes in *Argyroneta aquatica*. A. Phylogenetic analysis of ABC gene family of *A. aquatica* and other two spiders *Amaurobius ferox* and *Dolomedes plantarius.* Red line represent HGT-acquired ABC gene. B. Phylogenetic analysis of ACAD gene family of *A. aquatica* and other two spiders *A. ferox* and *D. plantarius.* Red line represent HGT-acquired ACAD gene. C. The features of the number of Exon and the GC counts of ABC and ACAD gene family among *A. aquatica, A. ferox* and *D. plantarius*. D. The exhibition of HGT genes in the scaffold 1 genome of *A. aquatica*. E. Collinearity analyses of some conserved genes near HGT-acquired genes in three species (*A. aquatica*, *A. ferox*, and *D. plantarius*) F. The heatmap of the HGT-acquired ABC and ACAD genes mapped to eight re-sequencing data of *A. aquatica* (P1-4: female; P5-8: male).

We pinpointed that these ABC and ACAD HGT-acquired genes were from bacteria, including 14 (29%) genes from Alcaligenaceae, 10 (20.5%) genes from Comamonadaceae, 10 (20.5%) genes from Moraxellaceae, 5 (10%) from Flavobacteriaceae, 4 (8%) from Rhodocyclaceae, 2 (4%) from Chitinophagaceae, and 4 (8%) from other family or order (Fig. S19A). And the main genus of the bacteria was *Acinetobacter*, *Alkanindiges*, and *Emticicia*. Notably, in the freshwater, evidence showed that all these bacteria can be found and confirmed (Fig. S19B).

### Comparative transcriptomics and metabolomics in water spider response to hypoxia stress

The water spider exhibits a remarkable ability to survive for over 48 hours with their spiracles closed using paraffin. We conducted comparative transcriptomics and metabolomics analyses to further investigate the molecular basis of hypoxia stress response in water spider.

Under hypoxia stress, comparative transcriptomic analysis of hypoxia group (LO) showed a total of 7 genes up-regulated and 17 genes down-regulated compared to control group (CK) (Fig. 4A). Among these genes, the significantly enriched functions were mainly related to energy metabolism, including *Gld* (Glucose dehydrogenase), *HAO1* (Hydroxyacid oxidase 1), *Enox1* (Ecto-NOX disulfide-thiol exchanger 1), *Slc13a5* (Solute carrier family 13 member 5), *AACS* (Acetoacetyl-CoA synthetase), and *SLC1A2* (Excitatory amino acid transporter 2). Notably, *HAO1* showed intensification selection, while *SLC1A2* showed relaxation selection.

**Figure 4.**
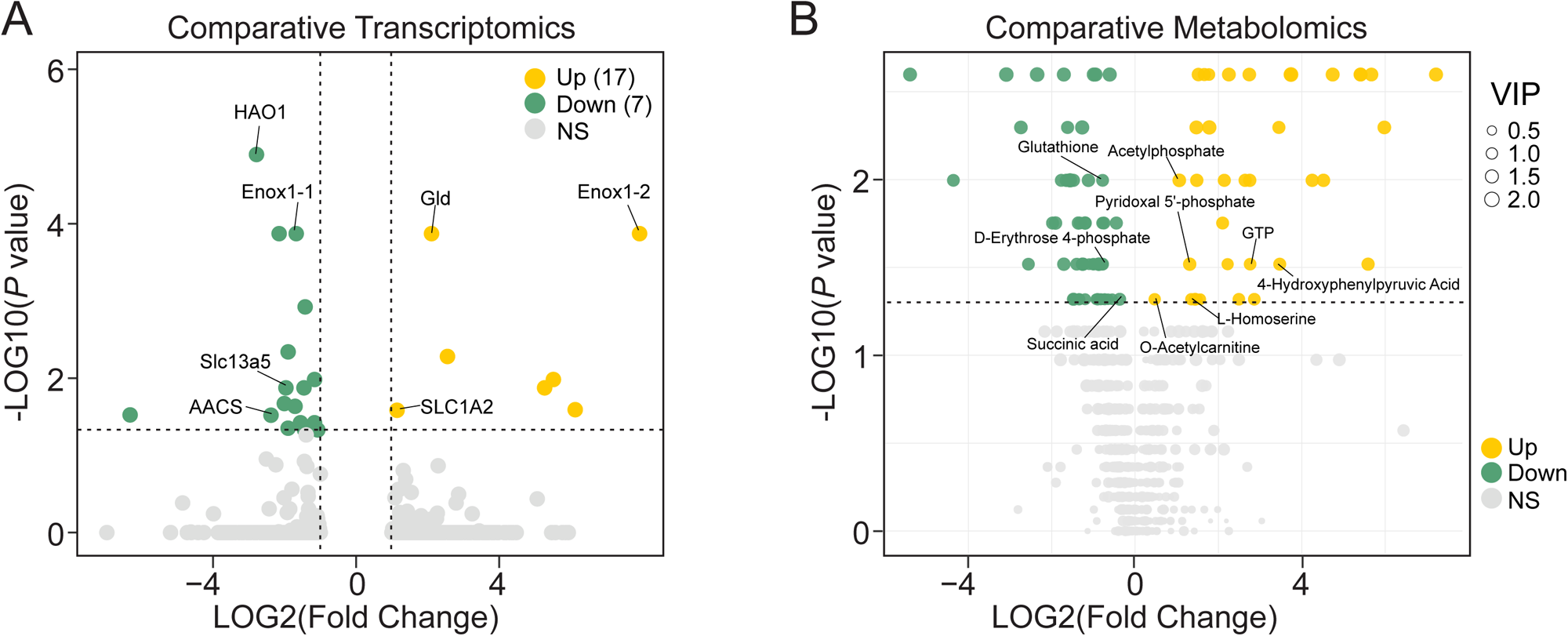
The compare transcriptome and metabonomics analysis of the hypoxia treatment group (LO) and the control group (CK). A. The volcano plots of the results of transcriptome for up- and down-regulation genes of CK vs LO groups. B. The volcano plots of the results of metabonomics for up- and down-regulation genes of CK vs LO groups.

By widely targeted metabolomics methods, in water spider, a total of 579 metabolites were identified and 95 differentiated metabolites (including 57 upregulated and 58 downregulated) were screened under hypoxia stress (Fig. 4B). Under hypoxia stress, the content of intermediate metabolites associated with major energy supply pathways such as glycolysis and TCA cycle changed significantly, for example, Pyridoxal 5’-phosphate, O-Acetylcarnitine, Acetylphosphate, L-Homoserine, GTP and 4-Hydroxyphenylpyruvic acid were up-regulation, while the D-Erythrose 4-phosphate, Glutathione, and Succinic acid were down-regulation in hypoxia stress treatment group, which involved in metabolism of glucide, lipid, protein and nucleic acid.

### Comparative tissue-specific and developmental-specific transcriptomes

Finally, we sought to investigate the spatial and temporal expression patterns of genes under selection or HGT genes in water spider. We performed multiple transcriptomic profiling of five tissues (venom, trachea, muscle, testis and ovary), and four development stages (3, 5, 6 and 19 days post fertilization). The expression patterns among different tissues or development stages of genes under relaxation, intensification and positive selection showed no significance (Fig. 5A, 5B).Interestingly, we observed most of HGT and non-HGT genes within ABC or ACAD gene family differentially expressed in all five tissues, while no HGT genes are expressed during the developmental stages.

**Figure 5.**
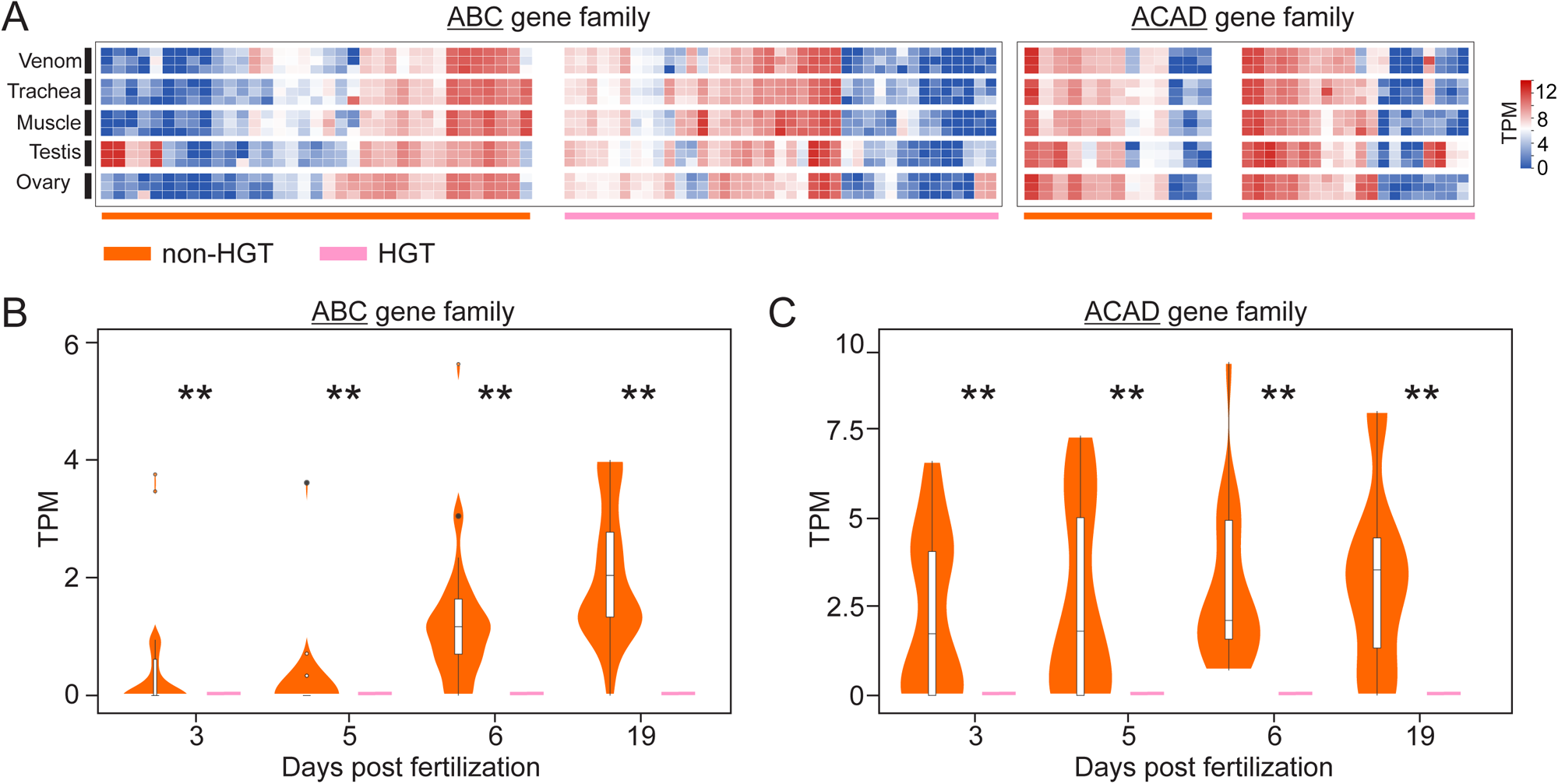
The expression of ABC and ACAD gene family among different tissues and develop stages. A. The heatmaps of non-HGT and HGT-acquired ABC and ACAD genes expression among different tissues including venom, trachea, muscle, testis and ovary. B. The expression of non-HGT and HGT ABC and ACAD gene family among 3, 5, 6 and 19 days post fertilization.

## Discussion

Understanding the genomic changes that enable spiders to adapt to underwater life is crucial, especially given their terrestrial ancestry. In this study, we conducted comparative genomic analyses of spiders from terrestrial, semi-aquatic, and aquatic habitats to address these questions. The marronoid spider clade includes species that inhabit a variety of specialized microhabitats, such as tundras, deserts, caves, intertidal areas, salt flats, and water, encompassing terrestrial, semi-aquatic, and aquatic environments (Gorneau et al., 2023; Leggett, Vink, and Nelson, 2024). Our study focused on the unique water spider *A. aquatica*, the semi-aquatic *Desis* genus, and various closely related terrestrial spiders within the marronoid clade. Notably, *A. aquatica* is the only spider that spends its entire life underwater. While previous studies have predominantly primarily explored the behavioral and physiological adaptations of water spiders to their aquatic lifestyle (Seymour and Hetz, 2011; Kang et al., 2014; Schaber et al., 2023), we provide a high-quality chromosome-level genome assembly of the water spider *A. aquatica*. To address the long-term lack of reference genomic resource for the semi-aquatic *Desis* genus and various closely related terrestrial spiders within in marronoid clade, we assembled and annotated the transcriptomic data of these spiders.

### Genomic evidence supports divergent evolutionary trajectories in semi-aquatic or aquatic adaptation

The evolution of semi-aquatic and aquatic spiders from terrestrial ancestors has been a topic of significant interest. Current taxonomy disperses these semi-aquatic and aquatic taxa across various families (Crews et al., 2020), suggesting that both types have independently evolved from terrestrial spiders (Leggett, Vink, and Nelson, 2024). Our comparative genomics analysis, integrated with life history and predation strategies, reveals distinct evolutionary routes for semi-aquatic and aquatic spiders adapting to underwater life. We observed a higher number of genes under intensified selection associated with the transition to aquatic life compared to semi-aquatic life. Specifically, the ratio of genes under intensified selection to those under relaxed selection was greater in the aquatic transition (92 intensification vs. 59 relaxation) than in the semi-aquatic transition (58 intensification vs. 67 relaxation). This relative abundance of intensified selection indicates that the evolutionary pressures to adapt to a fully aquatic environment are stronger and more urgent than those for a semi-aquatic environment. Adapting to an aquatic habitat requires comprehensive and often drastic genetic changes to survive and reproduce in water-dominated conditions, resulting in a higher number of genes under intensified selection. In contrast, the semi-aquatic transition involves organisms spending time both in water and on land. This dual lifestyle may not necessitate extensive genetic changes, or the selective pressures might be less intense because these organisms can still rely on terrestrial capabilities. Interestingly, there were more positively selected genes (PSGs) in semi-aquatic spiders (127 genes) compared to aquatic spiders (67 genes). This suggests that semi-aquatic spiders experience diverse and dynamic environmental pressures, requiring a greater number of genes to undergo positive selection during the adaptation to both terrestrial and aquatic environments. Aquatic spiders may have already undergone significant evolutionary adaptations to their environment, resulting in fewer genes currently under positive selection. Alternatively, the aquatic environment might present more stable or less varied selective pressures compared to the dual habitats of semi-aquatic spiders. Collectively, semi-aquatic spiders exhibit a higher number of positively selected genes due to the need to adapt to diverse conditions, while aquatic spiders display a higher number of genes under intensified selection, reflecting strong selective pressures for comprehensive adaptations to a fully aquatic environment.

### Respiratory adaptation in semi-aquatic and aquatic spiders

In aquatic and semi-aquatic environments, spiders encounter significant risks of respiratory distress and drowning, necessitating crucial adaptations to prevent asphyxia for their survival. Spider respiration depends on internal branching tubes and sacs connected to the atmosphere through spiracles on the abdominal wall (Foelix, 2011). These spiracles have a valvular mechanism that can remain open or partially closed to minimize water loss. Our study reveals significant advancements in the respiratory systems of semi-aquatic and aquatic spiders compared to their terrestrial counterparts. This evolution is particularly pronounced in fully aquatic spiders, which exhibit a highly refined gaseous exchange system (Fig. 2B). Among the genes with intensified or relaxed selection in water spiders, eight genes are associated with the development of the tracheae, as their functions have been reported in other animals. In water spiders, the *Wnt-1* and *Catenin beta* genes exhibited relaxed selection, with the Wnt/β-catenin pathway playing a pivotal role in trachea development (Llimargas and Lawrence, 2001; Gerhardt et al., 2018; Brechbuhl et al., 2011). The *TMEM234* gene shows intensified selection, while *TMEM59* exhibits relaxed selection. Although information regarding their structures, functions, and mechanisms within the TMEM family is limited, there is evidence suggesting that these gene family proteins are required for normal trachea development (Rock et al., 2008; Scholl et al., 2019; Sébastien et al., 2020). The *Arp3* and *ECE1* genes may regulate the diameter of the trachea and its contraction and expansion during respiration by participating in actin polymerization and contraction in smooth muscle (Zhang et al., 2005; De Campo et al., 2002). These genes also show intensified selection in water spiders. The *Syntaxin 5* gene, essential for the seamless growth and maintenance of tubular structures (Bourne et al., 2022), exhibits heightened selection pressure in water spiders. Additionally, *VEGF*, which controls cell migration during trachea development (Kipryushina et al., 2015), shows heightened selection pressure in water spiders. In semi-aquatic spiders, the gene *Hdac6* has been identified as under intensified selection and may function in the cilia of the trachea (Ran et al., 2016). This specific gene potentially plays a functional role in the respiratory adaptations required for the dual aquatic and terrestrial lifestyle of semi-aquatic spiders.

### Osmoregulatory adaptation in semi-aquatic and aquatic spiders

Osmoregulation is a critical physiological process for spiders living in aquatic or semi-aquatic environments. While cuticular structure and reduced metabolic rates help combat some osmotic stress, they are not sufficient on their own (Renault et al., 2016). Ion regulation is crucial for maintaining osmotic pressure. Under various experimental saline conditions, the concentrations of metal ions (Ca²⁺, K⁺, Mg²⁺, Na⁺) in spiders were reported to vary (Renault et al., 2016). The gene *Slc24a5*, which encodes a sodium/calcium/potassium exchanger involved in transmembrane ion transport, showed intensified positive selection in semi-aquatic spiders but exhibited relaxed selection in water spiders. Semi-aquatic spiders, such as those in the *Desis* genus, inhabit environments with high osmotic pressure, whereas water spiders face low osmotic pressure. In water spiders, horizontally acquired ABC genes, such as the high-affinity branched-chain amino acid transport ATP-binding proteins LivG and LivF, were identified. These genes provide the energy required to drive the transmembrane transport of branched-chain amino acids (Albers et al., 2004), potentially indirectly affecting osmotic pressure and regulation. Elevated salt loads may provoke DNA breaks, protein denaturation, and uncontrolled molecular interactions. Many genes related to DNA repair showed signs of intensified, relaxed, or positive selection among water and semi-aquatic spiders. In water spiders, genes such as *ALKBH2* and *PGBD4* exhibited relaxed selection, while *ppp4cb* showed intensified selection. In semi-aquatic spiders, genes such as *brcc3* and *Mes4* exhibited relaxed selection, while *Polr1b* and *Rad23*a showed intensified selection. Additionally, *KIN* and *Rad23a* genes exhibited positive selection. These findings underscore the complex and varied genetic adaptations that enable spiders to thrive in aquatic and semi-aquatic environments.

### Genetic adaptation in fat metabolism and digestion in semi-aquatic and aquatic spiders

In both aquatic and semi-aquatic spiders, various genes related to fat metabolism and digestion exhibit intensified or positive selection. For example, in semi-aquatic spiders, the genes *boca*, *ACADM*, *ABHD17B*, and *Acot13* show positive selection. The gene *Pnliprp2* exhibits both intensified and positive selection. In water spiders, the genes *Pnliprp2* and *PNLIP* display intensified selection, while *ND-ACP* shows positive selection. Additionally, many horizontally acquired ACAD genes were identified in water spiders, which are essential for energy production through mitochondrial fatty acid beta-oxidation (Shen et al., 2009). These genetic adaptations are particularly advantageous in environments where high-energy food sources, such as fish rich in long-chain fatty acids, constitute a primary diet (Nyffeler and Pusey, 2014; Shen et al., 2009).

### Hypoxia adaptation in aquatic spiders

Adaptation to hypoxia conditions is crucial for the successful transition of spiders from terrestrial back to aquatic environments. In water spiders, a variety of hypoxia-regulated genes involved in various aspects of energy metabolism have been identified as being under selection (Fuhrmann and Brune, 2017). Genes under intensified selection include *GPI*, *GFPT2*, and *UQCRC2*, while those under relaxed selection include *GlyP*, *slc37a3*, and *ETFA*. Under hypoxic conditions, enzyme reactions involving oxygen consumption may be influenced (Huwiler and Pfeilschifter, 2006). Notably, two genes encoding cytochrome P450, specifically *Cyp2j6* and *Cyp3a9*, have been identified as undergoing intensified selection. This finding aligns with the adaptive processes required for hypoxia tolerance in aquatic environments.

### Thermal adaptation in semi-aquatic Spiders

As poikilotherms, the semi-aquatic *Desis* spiders, being intertidal arthropods, require diverse strategies to cope with extreme temperature variations. The genes Heat shock factor-binding protein 1 (under relaxed selection) and *DNAJB4* (under positive selection) may play crucial roles in the cellular response to heat stress in these spiders.

In conclusion, these results provide critical insights into the genomic adaptations that enable spiders to thrive in underwater environments. By examining the roles of relaxed selection, intensified selection, positive selection, and HGT, we elucidate the genomic changes involved in the adaptation of spiders from terrestrial to semi-aquatic or aquatic habitats. Collectively, these findings not only enhance our understanding of the evolutionary paths of the semi-aquatic and aquatic spiders but also contribute to the broader knowledge of adaptive strategies in aquatic organisms.

## Methods

### Sample collection, sequencing and draft data collection

Water spider *A. aquatica* individuals were collected from Xilingol league of InnerMongolia (Figure 1A). Ten whole body of female adult water spiders without abdomen and stomach were used for whole genome sequencing. And an additional eight adults water spiders (four females and four males) were processed for whole genome resequencing. The tissues (including venom, muscle, trachea, testis, and ovary) and the embryo among different stages (3, 5, 6 and 19 days post fertilization) of water spider were used for transcriptome sequencing, each group contains three replicates. In addition, a female *Desis martensi* was collected from Sanya, Hainan Province (Figure 1A), followed by Starvation treatment for three days and used for transcriptome sequencing.

A combination of two sequencing approaches including short reads on MGISEQ 2000 platform and long reads on Oxford Nanapore Technology (ONT) PromethION platform (Nextomics Bioscience, Wuhan, China) had been performed for the whole genome sequencing. Specifically, the total genomic DNA for genome sequencing was extracted using Qiagen Blood & Cell Culture DNA Mini Kit. For short reads, the DNA libraries were prepared with the MGIEasy FS DNA Library Prep Set (BGI, Shenzhen, China) following the MGISEQ protocol, a total of 212.12 Gb from 2.13 billion clean reads were obtained. For long reads, the ONT libraries were prepared following the protocol of Ligation Sequencing Kit 1D (SQK-LSK108, ONT), the N50 of the raw nanopore reads length was 25 kb. An additional Hi-C sequencing was performed (Berry Genomics, Beijing, China). After cross-linked long-distance physical interactions and digested with MboI, the 300bp paired-end sequencing libraries were prepared for sequencing on an Illumina HiSeq PE150 platform. Finally, the transcriptome sequencing was performed on an Illumina NovaSeq 6000 platform (Berry Genomics, Beijing, China) yielding 150-bp paired-end reads. The details were shown in Table S11.

Besides the newly sequenced omics data, we download available genomes of terrestrial spider species including *Amaurobius ferox* (GCA_951213105.1), *Dolomedes plantarius* (GCA_907164885.2) and *Uloborus diversus* (GCF_026930045.1), and existing transcriptome data of spider species including a semi-aquatic spider *Desis japonica* (DRR296513), and terrestrial spiders *Metaltella simoni* (DRR296491), *Badumna longinqua* (DRR297554), *Adenodictyna kudoae* (DRR297439), *Dictyna felis* (DRR297156), *Brommella punctosparsa* (DRR296607), *Lathys maculosa* (DRR297435), *Cybaeus nipponicus* (DRR297040), *Cicurina japonica* (DRR297872), *Hahnia corticicola* (DRR296643), *Neoantistea quelpartensis* (DRR297193), *Agelena silvatica* (DRR297956), *Pireneitega luctuosa* (DRR297239), *Tegenaria domestica* (DRR297377), *Ambicodamus* sp. (DRR297731), *Pandava laminata* (DRR297807), *Nurscia albofasciata* (DRR296579) from NCBI Sequence Read Archive (https://www.ncbi.nlm.nih.gov/sra).

### Hypoxia stress treatment

For hypoxia treatment stress experiment, the lung slit and spiracle of adult male water spider was sealed by paraffin, then raised it in boxes at room temperature. After 48 hours, the spider was freezed with liquid nitrogen, removed the paraffin, and stored at −80 °C. The control group was not treated with paraffin seal, and raised it in boxes at room temperature. The details were shown in Figure S20. Finally, the samples were used for Liquid chromatography–mass spectrometry (LC-MC) assay and transcriptome sequencing (Suzhou PANOMIX Biomedical Tech, China). For LC-MC assay, each group contains eight replicates. For transcriptome sequencing, each group contains three replicates.

### The morphological analysis of respiratory systems

The four spiders including *Desis martensi*, *Badumna tangae*, water spider and *Dictyna felis* were used for morphological analysis of respiratory systems. All specimens were preserved in 75% ethanol in in the School of Life Sciences, Southwest University, Chongqing, China (SWUC). After washed by ddH2O, the specimen was then cleared by immersing them in pancreatin (Álvarez-Padilla and Hormiga 2007). The respiratory systems including book lungs and trachea were examined and measured using a Leica M205A stereomicroscope equipped with a drawing tube, a Leica DFC450 camera, and LAS software v4.6. Photos were taken with a Kuy Nice CCD mounted on an Olympus BX53 compound microscope. Compound focus images were generated using Helicon Focus 6.7.1 software.

### Genome assembly

The genome of water spider was assembled with one road of polishing iterations and the minimum overlap between reads of 3,000 using Flye v2.5 (Kolmogorov et al., 2019). The resulting genome assembly was further polished for two rounds using NGS data with NextPolish v1.0.5 (Hu et al., 2020). The Heterozygous regions were removed by Purge Haplotigs v1.1.0 (Roach et al., 2018) with the haplotigs identifying of 50% cut-off. The chromosome-level assembly was generated using Hi-C reads with Juicer v1.6.2 (Durand et al., 2016). To remove the potential contaminant sequences, the assembly genome was inspected against the NCBI nucleotide (nt) and UniVec databases by the software of HS BLASTN and BLAST+ (blastn) v2.7.1 (Chen et al., 2015). Finally, the quality and completeness of genome assemblies was assessed with BUSCO v5.4.5 (Simao et al., 2015) with arthropoda_odb10 data set (n= 1,013).

### Genome annotation

For the annotation of repetitive elements in genome, we searched it by means of a combination of ab initio and homology. Firstly, the species-specific repeat database (Ab-initio database) was constructed using RepeatModeler v2.0.1 (Flynn et al., 2020), and further combined the known repeat library (Repbase) and Ab-initio database as the reference repeat database. The repetitive elements in the genome was finally identified by RepeatMasker (Tarailo-Graovac and Chen, 2009).

Non-coding RNAs were annotated by Infernal v1.1.2 (Nawrocki and Eddy, 2013) and tRNAscan-SE v2.0.6 (Chan et al., 2021). In addition, a tool “EukHighConfidenceFilter” in tRNAscanSE was used for further filtering transfer RNAs (tRNAs) with high confidence.

For the annotation of gene structure, we integrated the ab initio, transcriptome-based, and protein homology-based evidence by Maker v2.31.10 (Holt and Yandell, 2011). Gene structure prediction of ab inito was identified by Augustus v3.3.3 (Hoff and Stanke, 2019) and GeneMark-ES/ET/EP v4.48 3.60 lic (Bruna et al., 2020). To model the sequence properties accurately, both gene finders (Augustus and GeneMark-ES/ET/EP) were initially trained using the BRAKER v2.1.5 pipline (Bruna et al., 2021). The transcriptome data for whole body water spider was mapped to the genome by the software HISAT2 v2.2.0 (Kim et al., 2015). BRAKER pipeline was run with default parameters. Then further assembled RNA-Seq alignments into potential transcripts by Stringtie v2.1.3 (Pertea et al., 2015), and the results was provided as input for Maker via the “est” option. The protein homology-based evidence was download from NCBI database, which including the protein sequences of *Stegodyphus mimosarum* (GCA 000611955.2), *Parasteatoda tepidariorum* (GCA 000365465.3), *Trichonephila clavipes* (GCA 002102615.1), *Drosophila melanogaster* (GCA 000001215.4), *Ixodes scapularis* (GCA 002892825.2), *Daphnia pulex* (GCA 900092285.2) and *Strigamia maritima* (GCA 000239455.1), which were combined and fed to Maker pipline.

Gene function annotation was predicted by Diamond v0.9.24 (Buchfink et al., 2015), InterProScan v5.41–78.0 (Mulder and Apweiler, 2007) and eggNOG-mapper v2.0 (Cantalapiedra et al., 2021). The UniProtKB/Swissprot database was against by the software Diamond with the mode of more sensitive, maximum number for target sequences of 1, and e-value of 1e-5. The following 5 databases including Pfam (Mistry et al., 2021), Panther (Mi et al., 2016), Gene3D (Lewis et al., 2018), Superfamily (Pandurangan et al., 2019), and Conserved Domain Database (CDD) (Marchler-Bauer et al., 2017) was against by the software InterProScan. The the eggNOG v5.0 database (Huerta-Cepas et al., 2019) was against by the software eggNOG-mapper.

Due to the annotation files for *A. ferox* and *D. plantarius* are not publicly available, we also performed separate annotations for these two species in this project.

### RNA-seq assembly and annotation

We assembled and annotated the transcriptomes data in these study. Quality control of the raw data was performed using BBTools suite v38.67 of “bbduk.sh” for trimming the reads’ ends to Q20 with reads shorter than 15 bp or with >5 Ns. Then the transcript sequences was assembled with de novo assembly using Trinity v.25.1 (Grabherr et al., 2011). The redundancy sequences were removed by cd-hit-est v4.8.1. And the assembled transcript sequences were annotated by Trinotate v3.0.1 (Bryant et al., 2017).

### Species phylogeny and gene orthology

A total of eight families in marronoid spider clade including Amaurobiidae (*A. ferox*), Agelenidae (*A. silvatica*, *P. luctuosa*, *T. domestica*), Desidae (*D. japonica*, *D. martensi*, *M. simoni*, *B. longinqua*), Hahniidae (*H. corticicola*, *N. quelpartensis*), Cybaeidae (*C. nipponicus*), Cicurinidae (*C. japonica*), Dictynidae (*A. kudoae*, *D. felis*, *L. maculosa*), and five closely related families spidcies including Cicurinidae (*B. punctosparsa*), Nicodamidae (*Ambicodamus* sp.), Titanoecidae (*P. laminata*, *N. albofasciata*), Pisauridae (*D. plantarius*), and the Uloboridae (*U. diversus*) as outgroup were combined with water spider *A. aquatica* for Gene clusters analysis. The proteins of these species were fed to OrthoFinder v2.5.4 (Emms et al., 2019) for orthologous gene clusters analysis.

For phylogenetic analysis, a total of 166 single copy orthologous genes were used to examine the phylogenetic relationships. First, the genes were separately aligned by MAFFT v7.471 (Katoh and Standley, 2013) with L-INS-I strategy. The resulting alignments were then trimmed by trimAl v1.4 (Capella-Gutierrez et al., 2009) with method “automated1”. Then the results were concatenated by FASconCATG v1.04 (Kuck and Longo, 2014). Finally, the maximum likelihood (ML) tree was constructed by IQ-TREE v2.0.3 (Minh et al., 2020) with the optimization number of rcluster algorithm, replicates for ultrafast bootstrap, and replicates for Shimodaira–Hasegawa (SH) approximate likelihood ratio tests being 10, 1000, and 1000, respectively. The divergence time calibration was performed with the reference from Magalhaes (Magalhaes et al., 2020; Moradmand, Schonhofer and Jager, 2014), paleobiodb database (https://paleobiodb.org/) and TimeTree database (http://www.timetree.org/), with Hahniinae (43-47.8 Mya), Dictynidae (100-189.7 Mya), and Entelegynae (132.9-140 Mya). The divergence time was calculated using package MCMC Tree in PAML v4.10.0 (Yang, 2007).

### Pattern of molecular evolution in shared orthologs

To determine the patterns of molecular evolution of terrestrial, semi-aquatic and aquatic spiders in the set of semi-aquatic and aquatic branches across the phylogeny, we characterized the rates of nonsynonymous to synonymous rate (dN/dS) in each shared ortholog. The protein sequence alignment was performed by MAFFT v7.471 (Katoh and Standley, 2013). Then the codon alignments of these orthologs which derived from amino acid alignments and corresponding DNA sequences using PAL2NAL v.14 (Suyama et al., 2006). And the shared ortholog alignments with length of at least 50 codons was executed filtration. The discrete codon substitution rate (dN/dS, ω) representing water spider, semi-aquatic spider and terrestrial spider species were estimated by HyPhy (Pond et al., 2005) following the pipeline as previously described (Tong et al., 2022).

Further, the pattern of selection acting on the semi-aquatic transition and aquatic transitionfrom was detected by RELAX (Wertheim et al., 2015) at gene-wide level. Genes are considered to experience significantly relaxed selection with a K-value < 1 (q-value < 0.05). Conversely, when K-value > 1 (q-value < 0.05), the genes are considered to experience significantly intensified selection. The q-values represent an estimate of the false discovery rate (FDR) for each gene which were calculated using the Benjamini–Hochberg method.

Finally, the signal of positive selection in genes involved in semi-aquatic and aquatic transitions were detected by BUSTED-PH (https://github.com/veg/hyphy-analyses/blob/master/BUSTED-PH/README.md) (Murrell et al., 2015) and followed by the correcting for multiple testing (Benjamini–Hochberg). We defined the genes under positive selection with q-value < 0.05 for foreground branches (i.e. aquatic spider), q-value > 0.05 for background branches (i.e. territorial spiders), and q-value < 0.05 for the difference between foreground and background branches.

### HGT gene identification and validation

To determine if specific genes in water spider that may have been horizontally acquired, we employed conservative phylogeny-based approach following the methodology (Li et al., 2022). For ABC and ACAD genes in water spiders, we first blast these genes with corresponding protein sequence by BLASTP in NCBI database.

To further validate the putative HGT-acquired genes, we compared them against previously published genes across the phylogenetic tree. We then analyzed gene expression across different tissues to assess their biological relevance. Additionally, we selected nine putative HGT-acquired genes—six ABC genes and three ACAD genes—for further experimental validation. For these genes, we performed PCR assays to amplify regions flanking the foreign gene, targeting sequences between 700 and 1,500 bp. The PCR products were analyzed via agarose gel electrophoresis to confirm the expected sizes, and successful amplifications were sequenced using Sanger sequencing. If the flanking regions of the HGT-acquired genes were successfully amplified and the Sanger sequences showed over 98% identity with the reference genome, the gene was considered validated. Primer sequences for these PCR assays are provided in Table S12. Additionally, we visualized the distribution of HGT-acquired genes on scaffold 1 of the water spider genome using the online tool MG2C v2.1 (Chao et al., 2021).

To confirm the HGT-acquired genes available in genome, the kallisto v0.48.0 (Bray et al., 2016) was used for counts the reads for HGT-acquired genes among the six water spider re-sequencing genome reads.

### Differential gene expression analysis

To explore the changes in gene expression response to hypoxia stress, we performed comparative transcriptomic analysis between spider samples under hypoxia stress and normal oxygene condition as control. For this, we aligned the transcriptome sequencing reads of whole body to the whole genome of water spider by HISAT2. The read counts were counted by featureCounts v1.6.4 (Liao et al., 2014) and then normalized among samples FPKM normalization method by DESeq2 package v1.24.0 (Love et al., 2014). Genes with an adjusted p value <0.05 and |log2 ratio| ≥ 1 were considered to be differentially expressed. The volcano plot was generated using the “ggplot2” package in R.

### Differential metabolites analysis

To investigate differential metabolites induced by hypoxia treatment, we conducted non-targeted metabolomics using LC-MS (Suzhou PANOMIX Biomedical Tech, China) to compare spider samples under hypoxic and normal oxygen conditions. The sample preparation followed these steps: first, 1g of sample was accurately weighed and mixed with 1000 µL of tissue extract (75% methanol [9:1], 25% H2O) and three steel balls in a 2mL centrifuge tube. The samples were then ground in a tissue grinder at 50 Hz for 60 seconds, repeating the process twice. After grinding, the samples underwent ultrasonic shaking for 30 minutes, followed by an ice bath for 30 minutes. The mixture was centrifuged at 12,000 rpm for 10 minutes at 4°C, and the supernatant was collected and dried. To prepare the samples for LC-MS detection, 200 µL of 50% acetonitrile solution containing 2-Amino-3-(2-chlorophenyl)-propionic acid (4 ppm) was used to re-dissolve the dried samples. The supernatant was filtered through a 0.22 μm membrane and transferred to a detection vial for LC-MS analysis. Various statistical methods, including principal component analysis, clustering, and univariate analysis, were applied to compare metabolite profiles between the different groups. The volcano plot illustrating significant metabolite changes was generated using the “ggplot2” package in R.

## Data availability

The raw Nanopore data are available on NCBI with the project of PRJNA559680 with the SRA accession numbers of SRR9994193. And the whole-body RNA accession number is SRR9994194. For water spider *Argyroneta aquatica,* the genome sequence (https://cstr.cn/31253.11.sciencedb.14858), the raw Illumina data (https://cstr.cn/31253.11.sciencedb.11012), the Hi-C data (https://doi.org/10.57760/sciencedb.15431), the metabolites and transcriptomic raw data (https://cstr.cn/31253.11.sciencedb.15069), the RNA data for different stage of embryo (https://cstr.cn/31253.11.sciencedb.10923), and the RNA data of different tissues (https://cstr.cn/31253.11.sciencedb.15029) are available on scienceDB. And the RNA data of a female *Desis martensi* (https://cstr.cn/31253.11.sciencedb.09032 is also available on scienceDB.

## Acknowledgements

We thanked Kun Yu and Feng Lu for provide photos of water spider. This work was supported by National Key R&D Program of China (Grant No. 2022YFC2601200, 2023YFC2604904), the National Science & Technology Fundamental Resources Investigation Program of China (Nos. 2023FY100301,2022FY100500), National Science Foundation of China (Nos. 32270468,32261143728), the project of the Northeast Asia Biodiversity Research Center (NABRI202203) by Ming Bai. This research is also supported by the Science Foundation of School of Life Sciences SWU (20232008071901 and 20212020110501) and the Natural Science Foundation of Chongqing (No. cstc2019jcyj-zdxmX0006) by Zhi-Sheng Zhang.

## Author contributions

C.T., M.B. and Z.Z. designed the original research. Z.F. and C.T. performed data analysis and drafted the manuscript. B.L., L.S., T.R. and P.L. performed sample preparation and data analysis. L.W., J.G., L.C., and B.T. collected the samples and performed sample preparation. Y.C. drew the illustration. Q.H., M.D., Q.Z., X.Z., J.L., N.L. and M.I. performed formal analyses and visualization of the data. All authors revised and approved the final manuscript.

