## Supplemental Figure 1-20 for "Genomic adaptations to semi-aquatic and aquatic life in spiders"

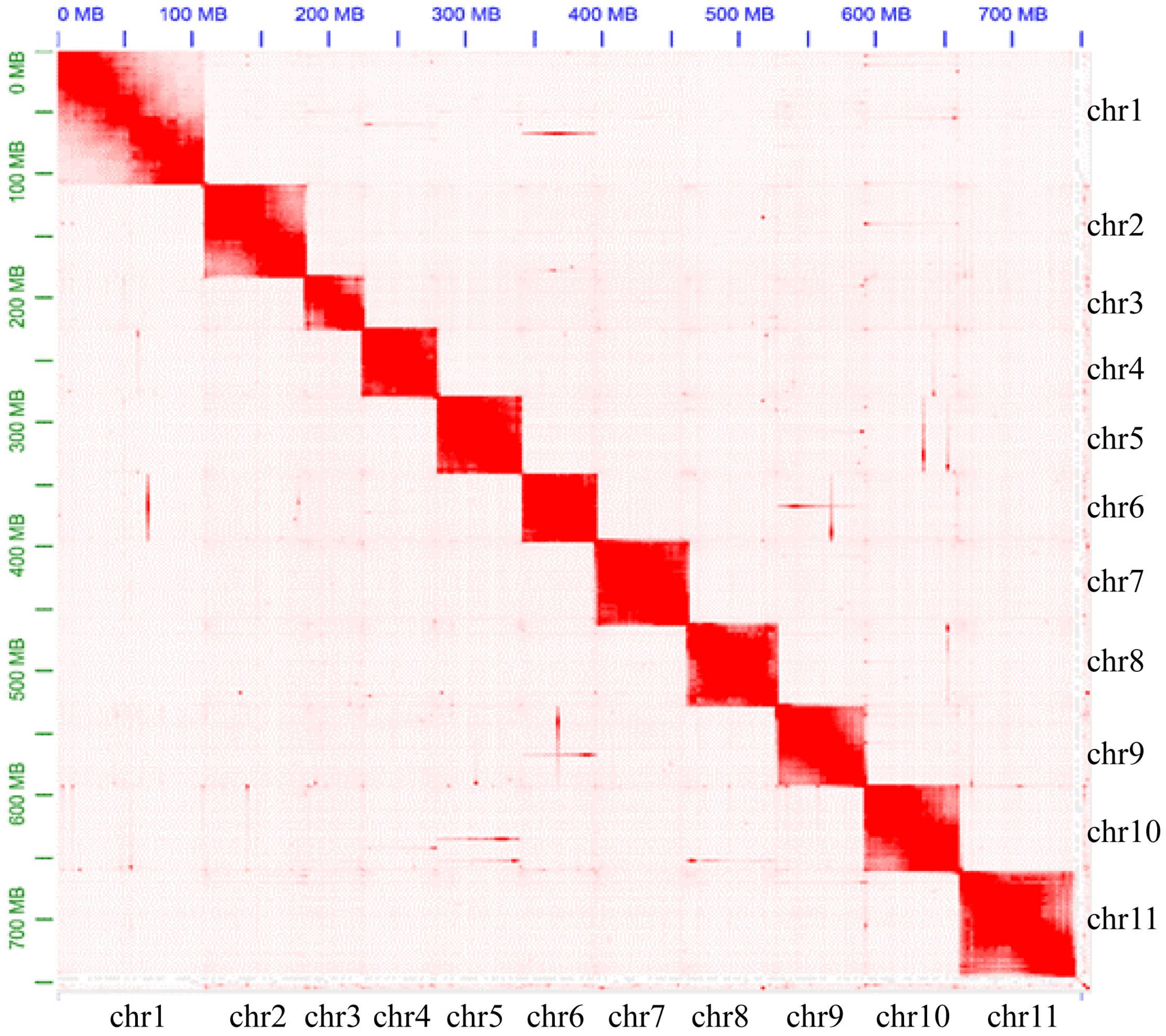


Figure S1. Hi-C interaction heat map for water spider genome showing interactions among 11 chromosomes.

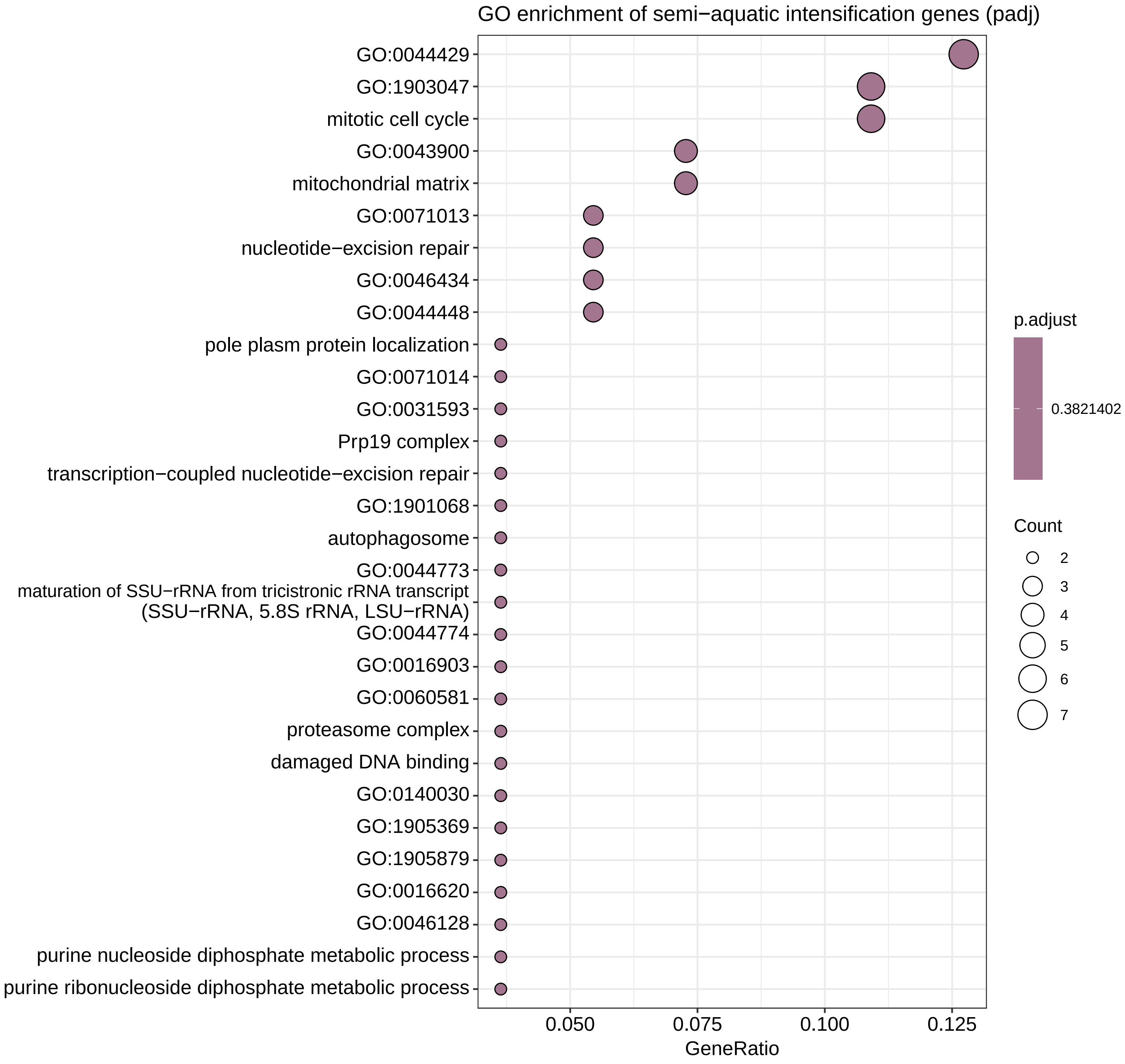


Figure S2. GO enrichment of semi-aquatic intensification genes.

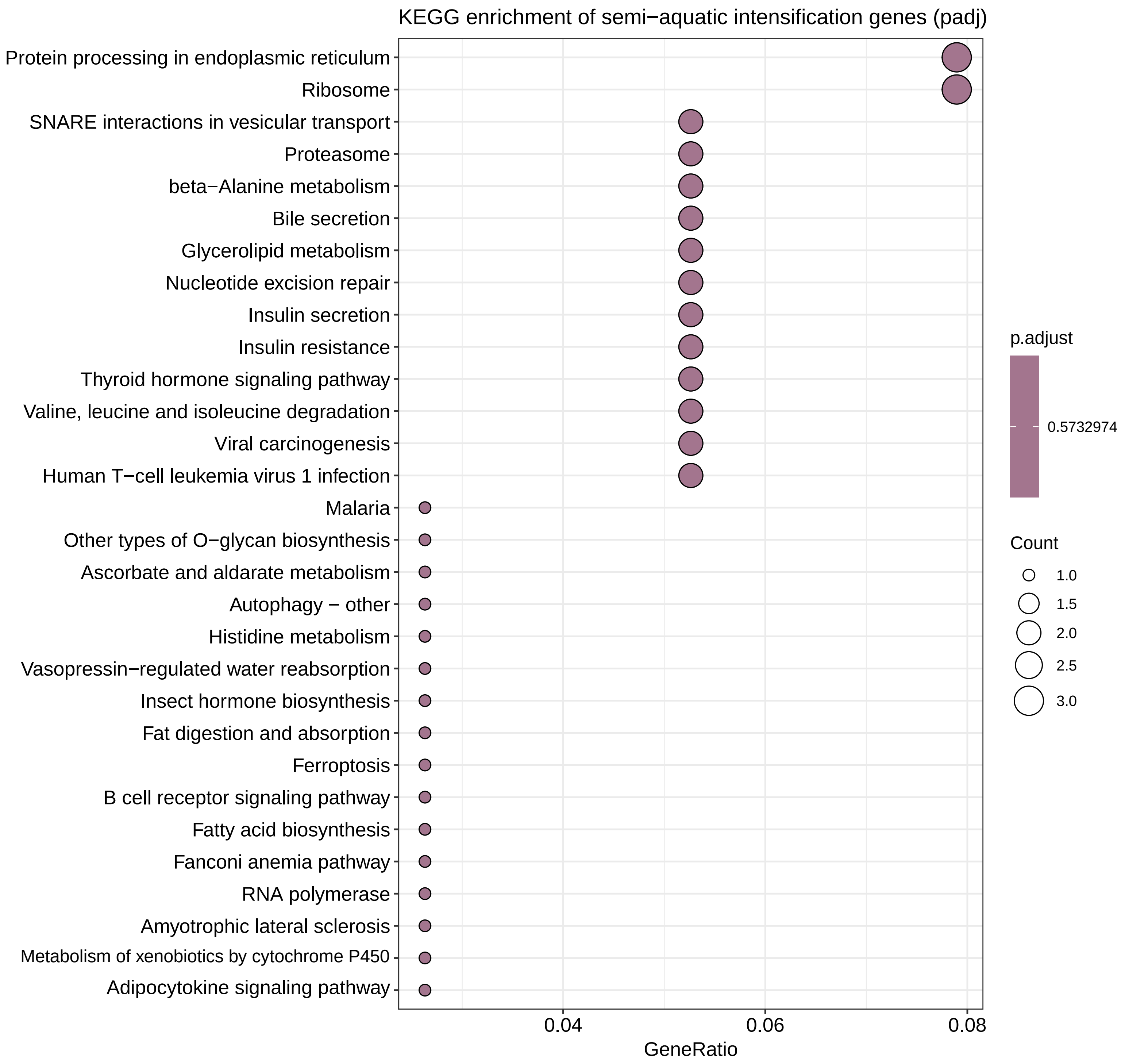


**Figure S3. KEGG enrichment of semi-aquatic intensification genes.**

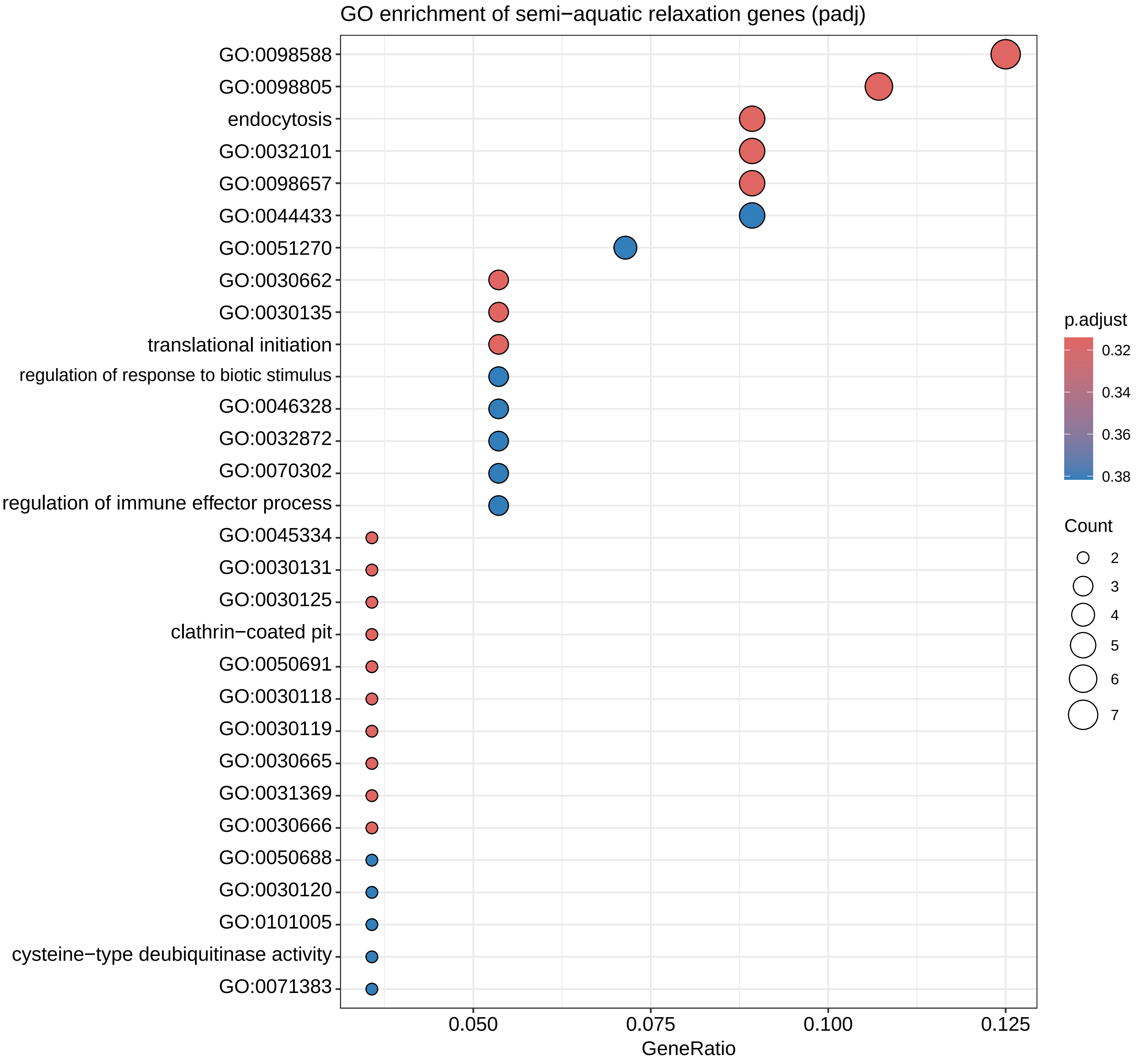


**Figure S4. GO enrichment of semi-aquatic relaxation genes.**

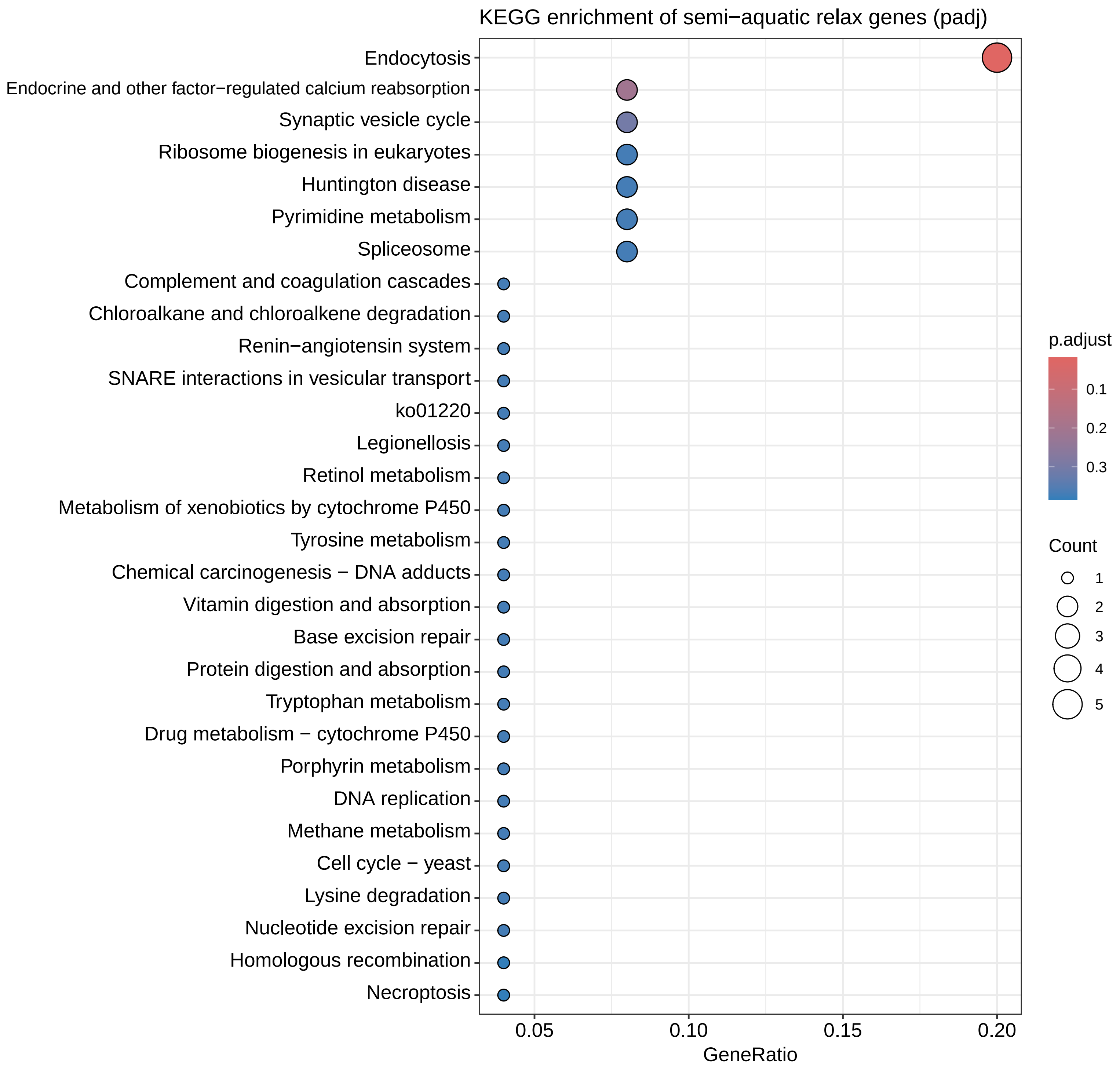


**Figure S5. KEGG enrichment of semi-aquatic relaxation genes.**

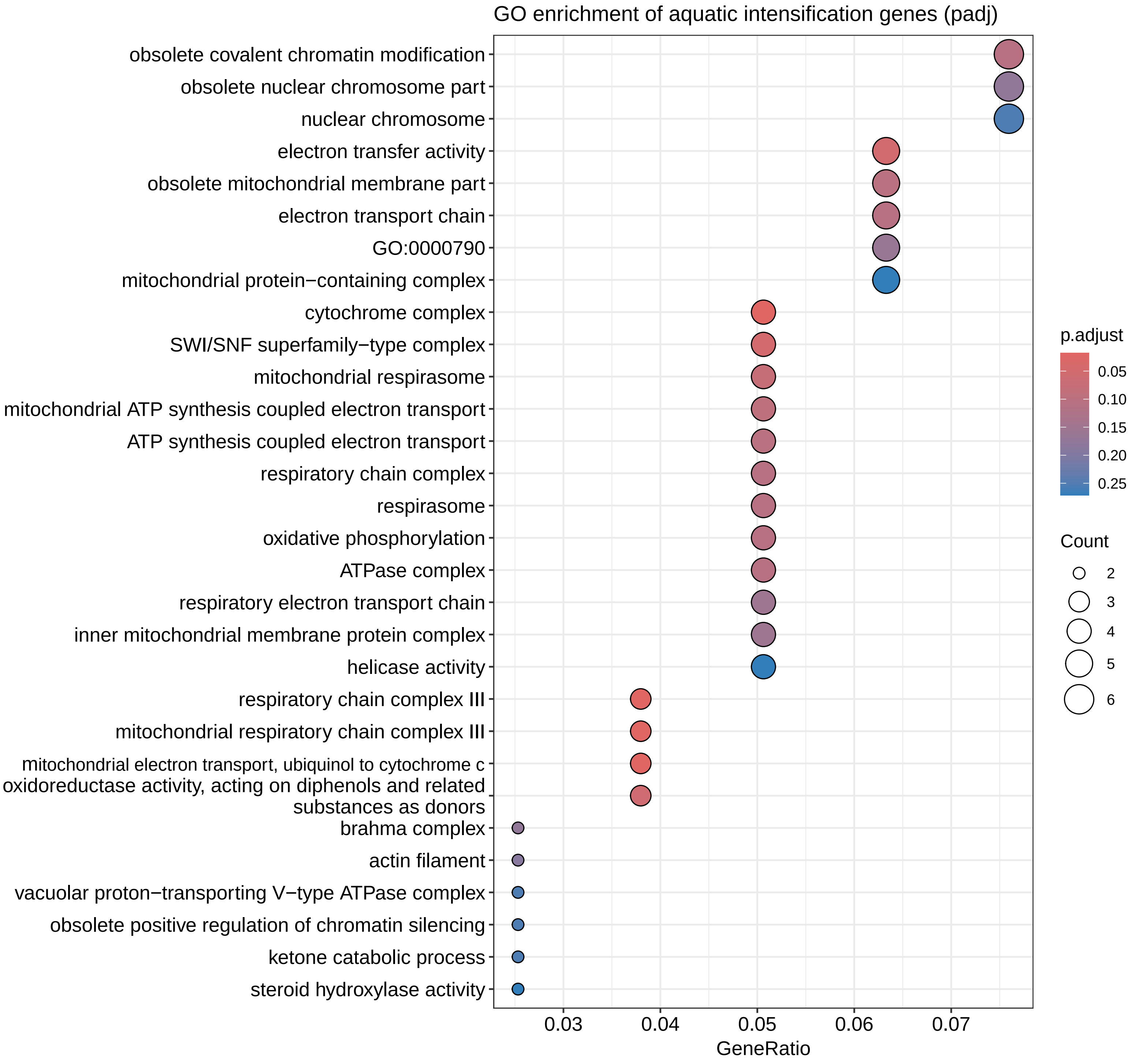


**Figure S6. GO enrichment of aquatic intensification genes.**

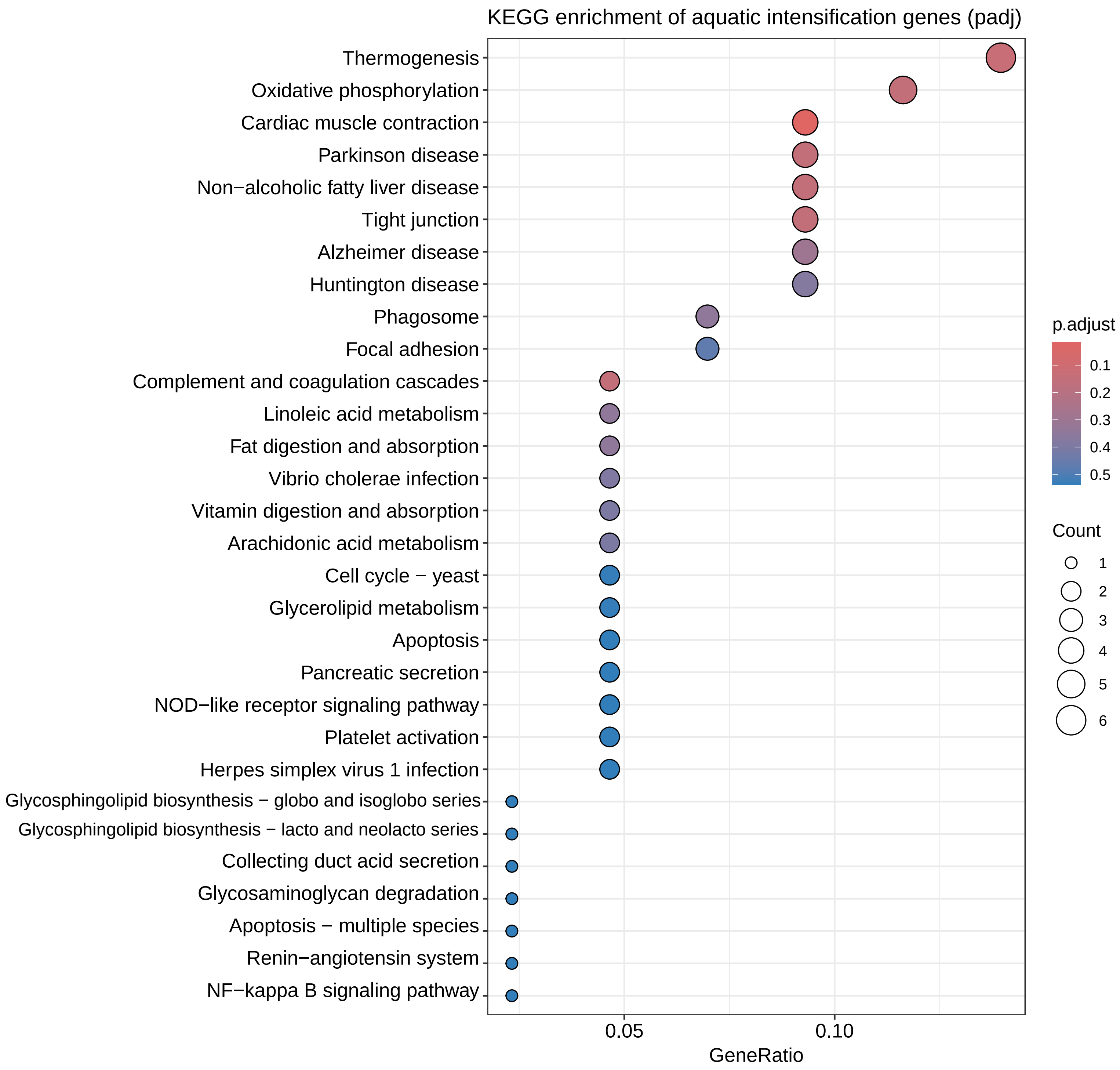


**Figure S7. KEGG enrichment of aquatic intensification genes.**

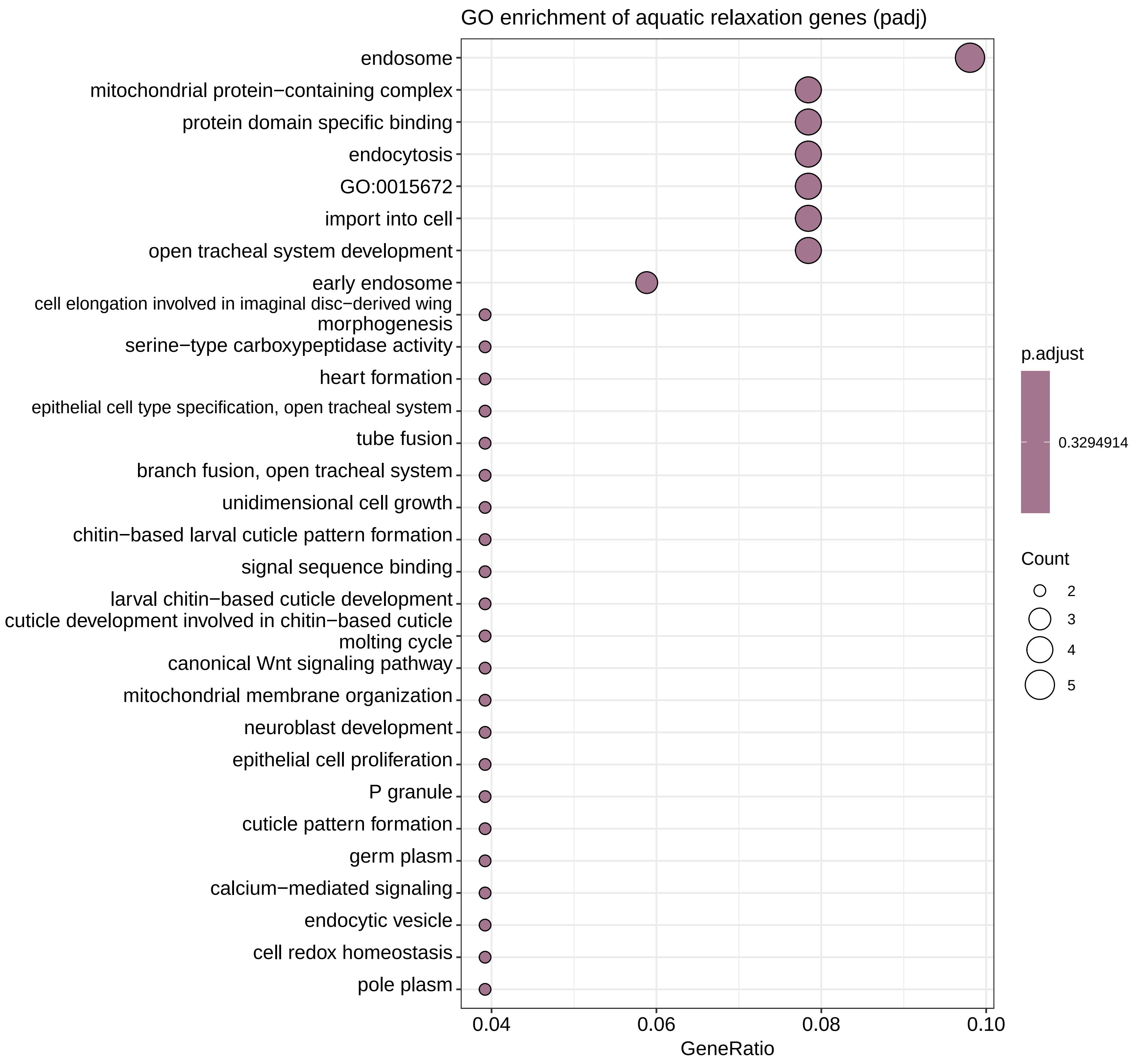


**Figure S8. GO enrichment of aquatic relaxation genes.**

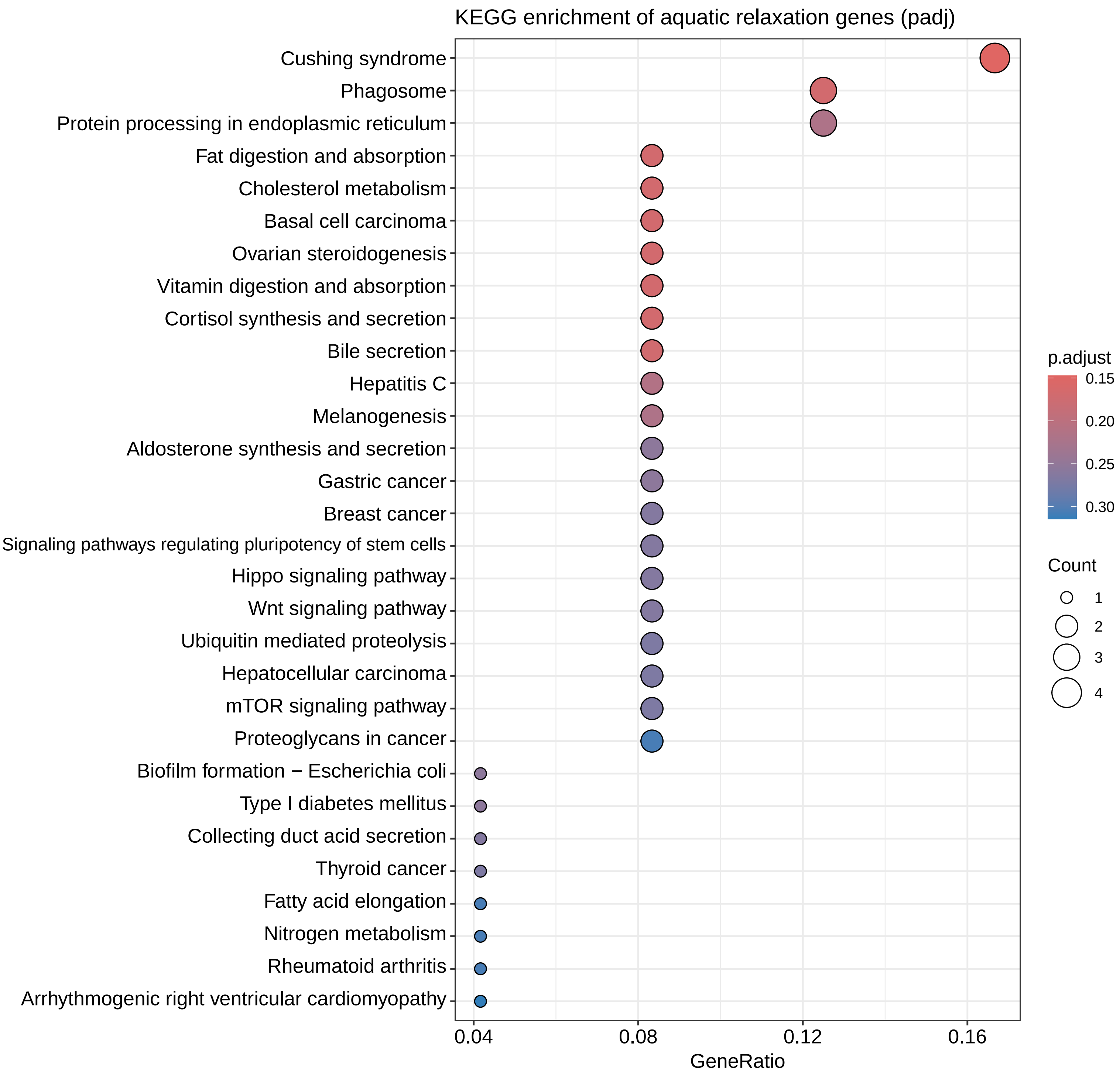


**Figure S9. KEGG enrichment of aquatic relaxation genes.**

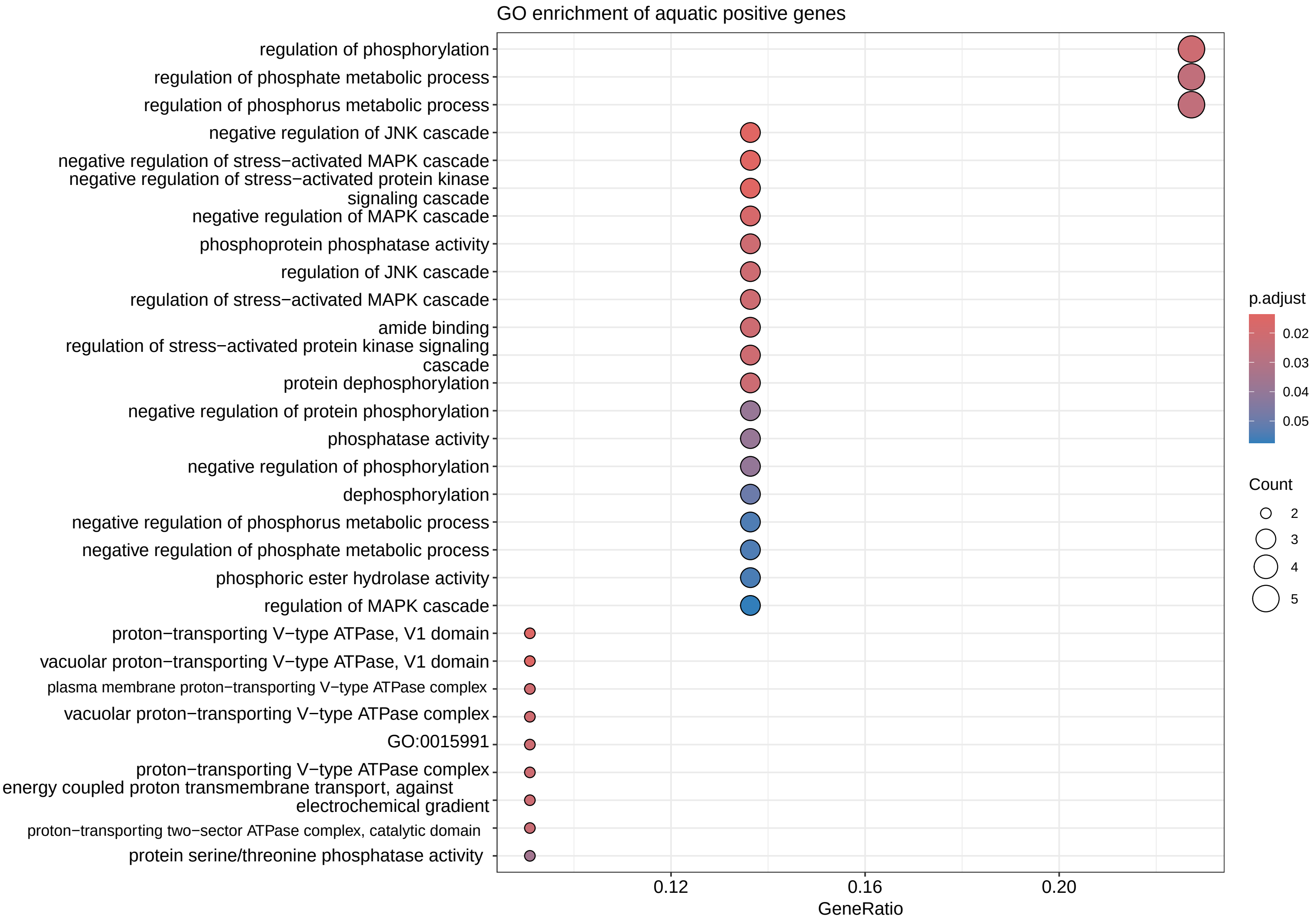


**Figure S10. GO enrichment of aquatic positive genes.**

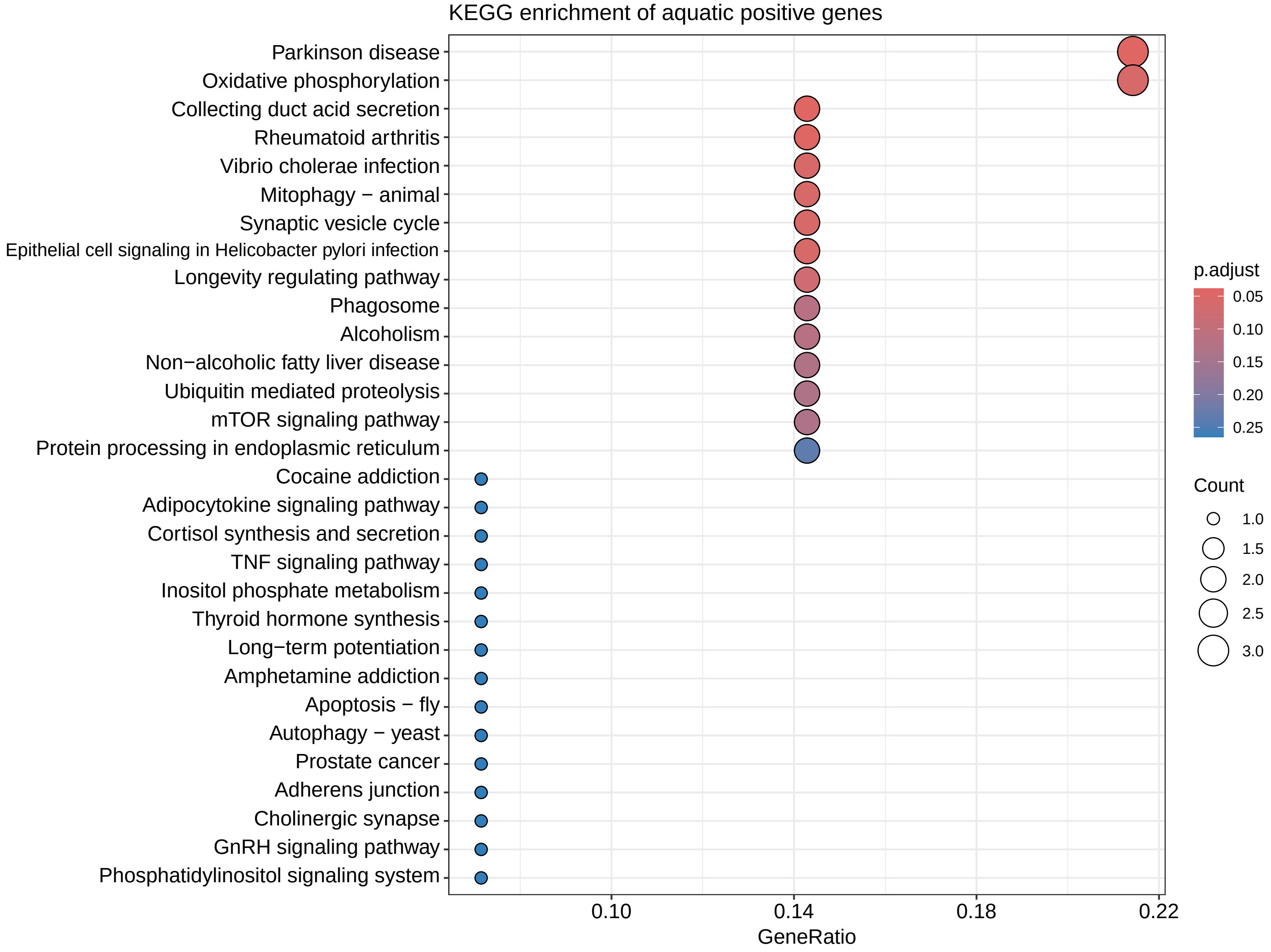


**Figure S11. KEGG enrichment of aquatic positive genes.**

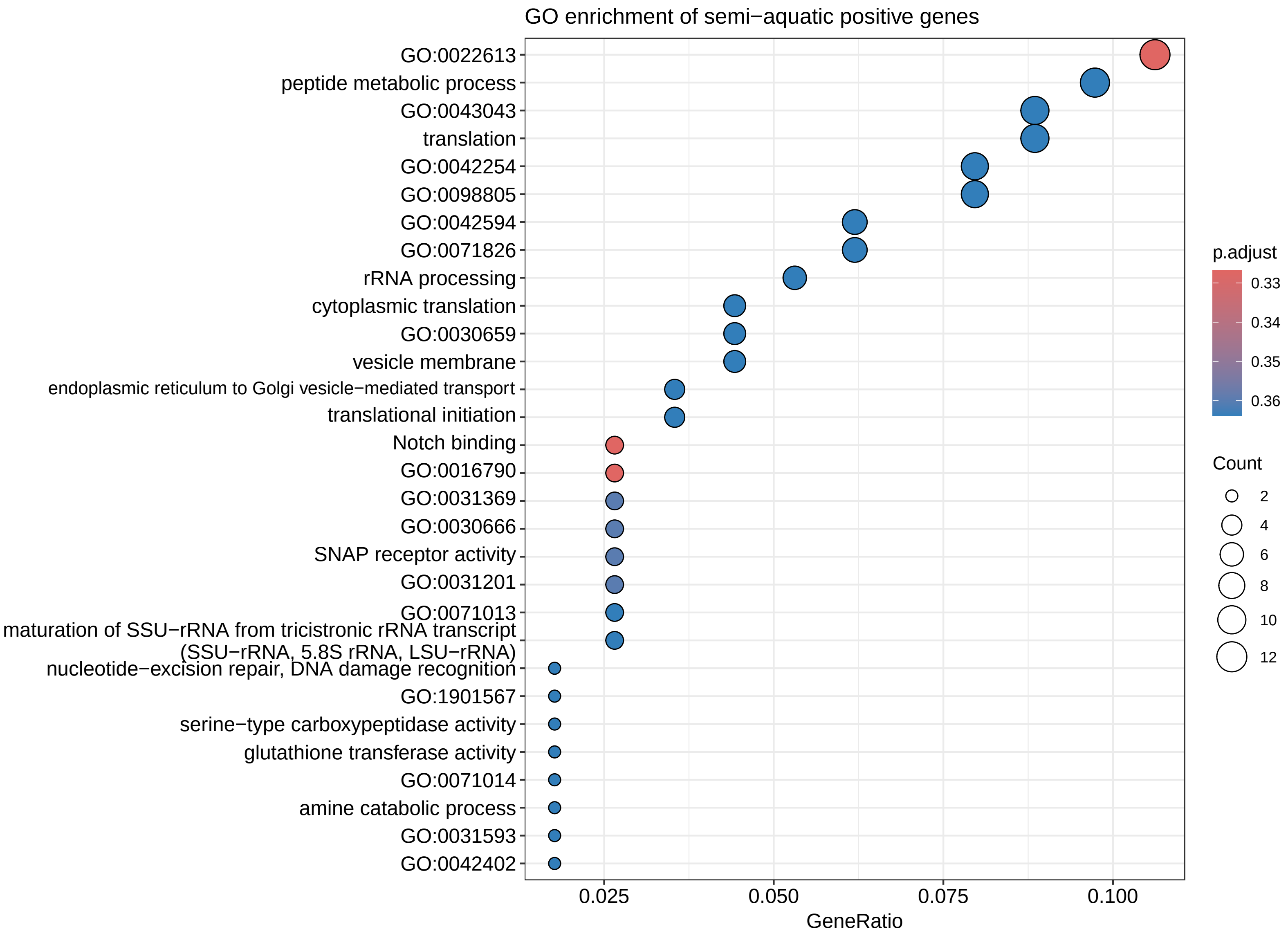


**Figure S12. GO enrichment of semi-aquatic positive genes.**

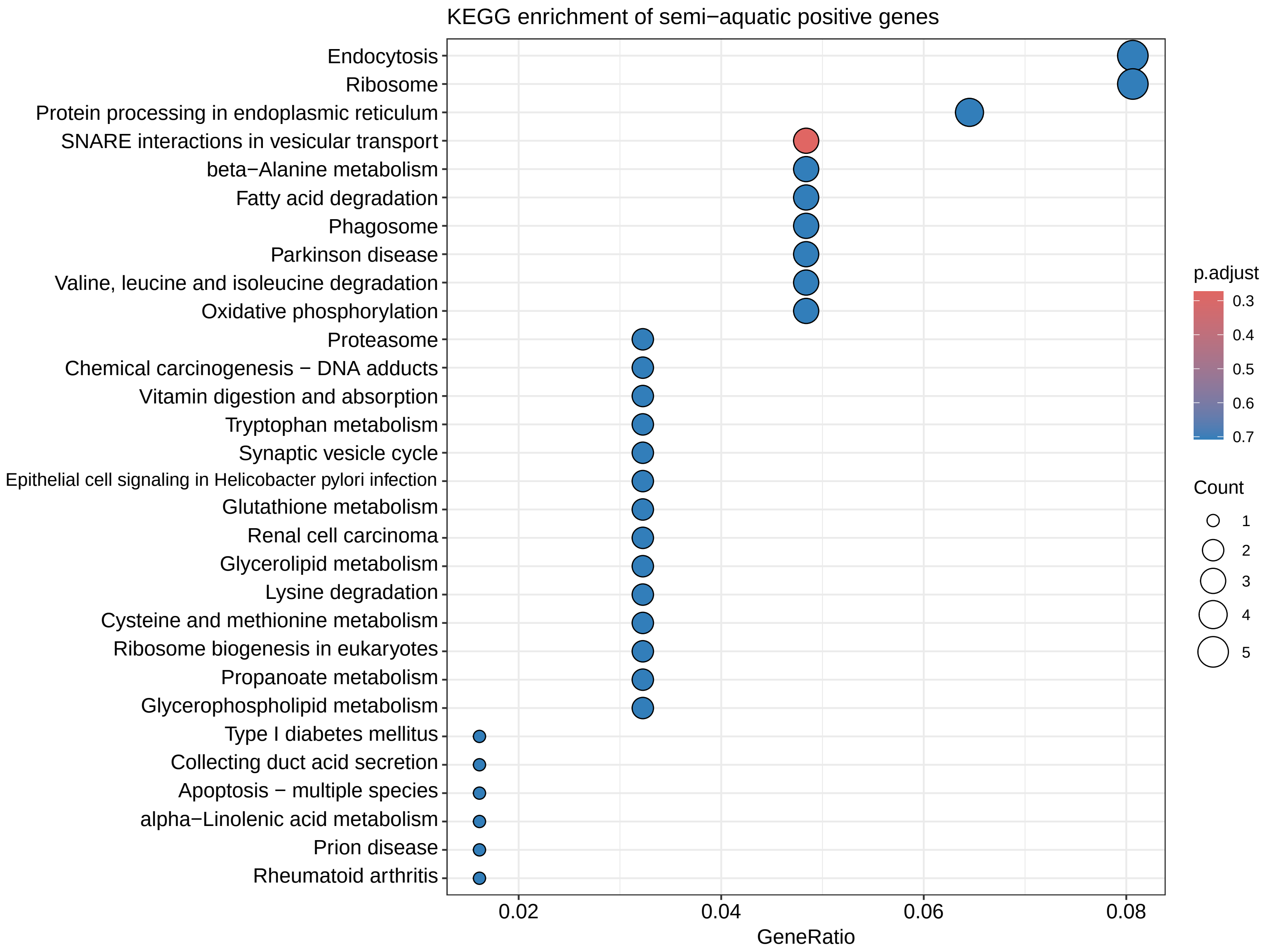


**Figure S13. KEGG enrichment of semi-aquatic positive genes.**

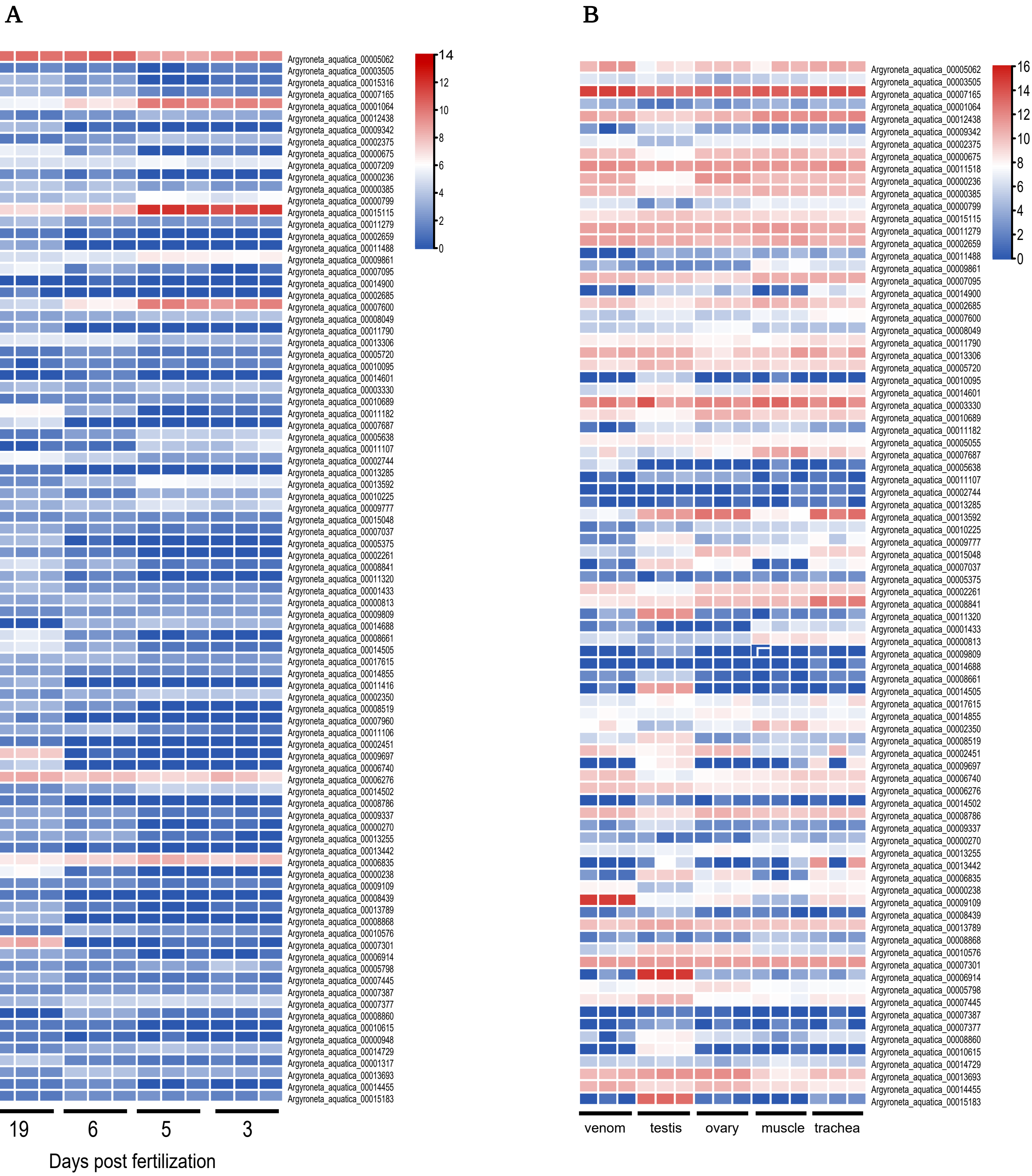


**Figure S14. The expression of aquatic intensification selection genes among different tissues and develop stages.**
A. The expression of aquatic intensification genes among 3, 5, 6 and 19 days post fertilization.
B. The heatmaps of aquatic intensification genes expression among different tissues including venom, trachea, muscle , testis and ovary.

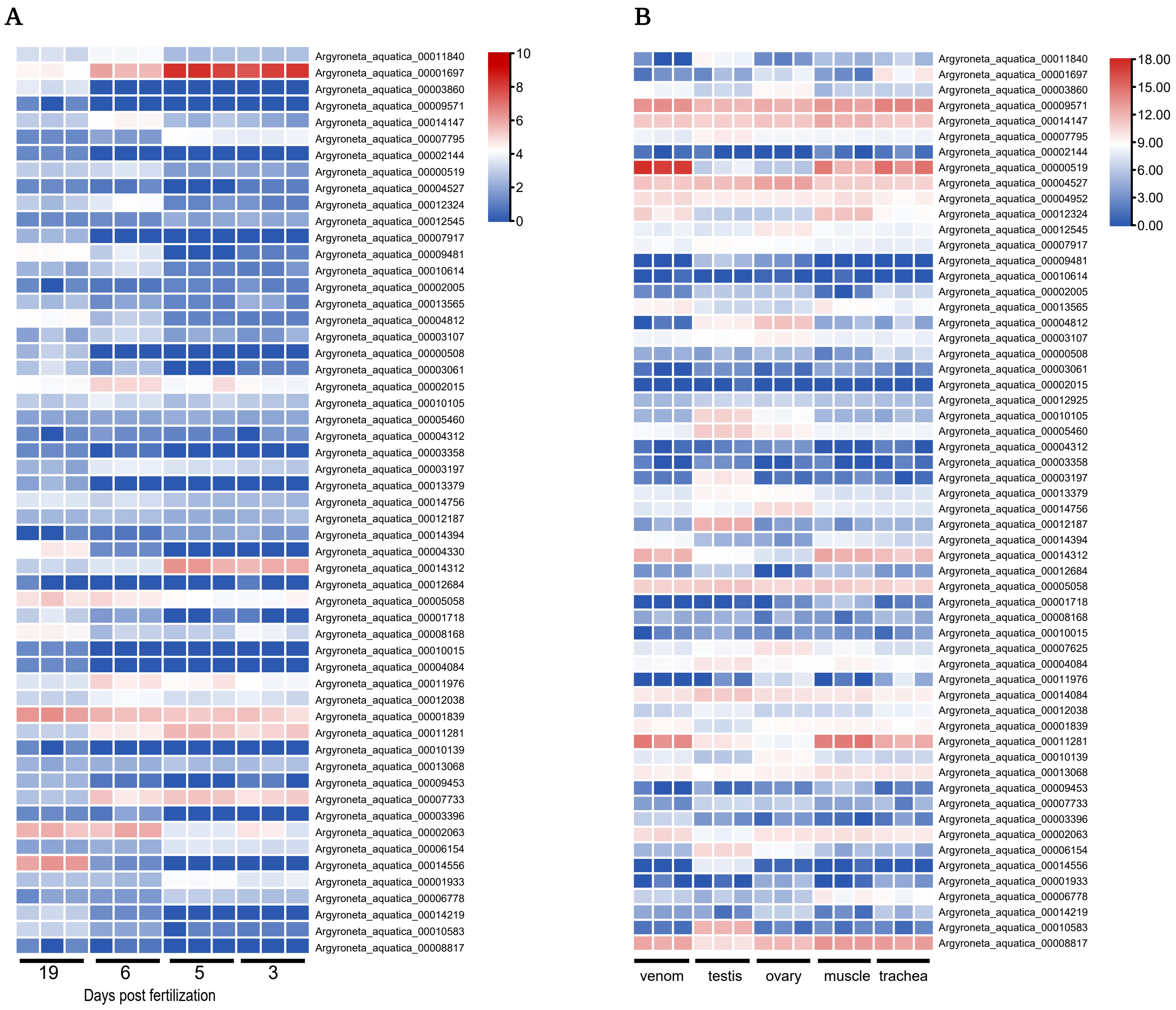


**Figure S15. The expression of aquatic relaxation selection genes among different tissues and develop stages.**
A. The expression of aquatic relaxation genes among 3, 5, 6 and 19 days post fertilization.
B. The heatmaps of aquatic relaxation genes expression among different tissues including venom, trachea, muscle, testis and ovary.

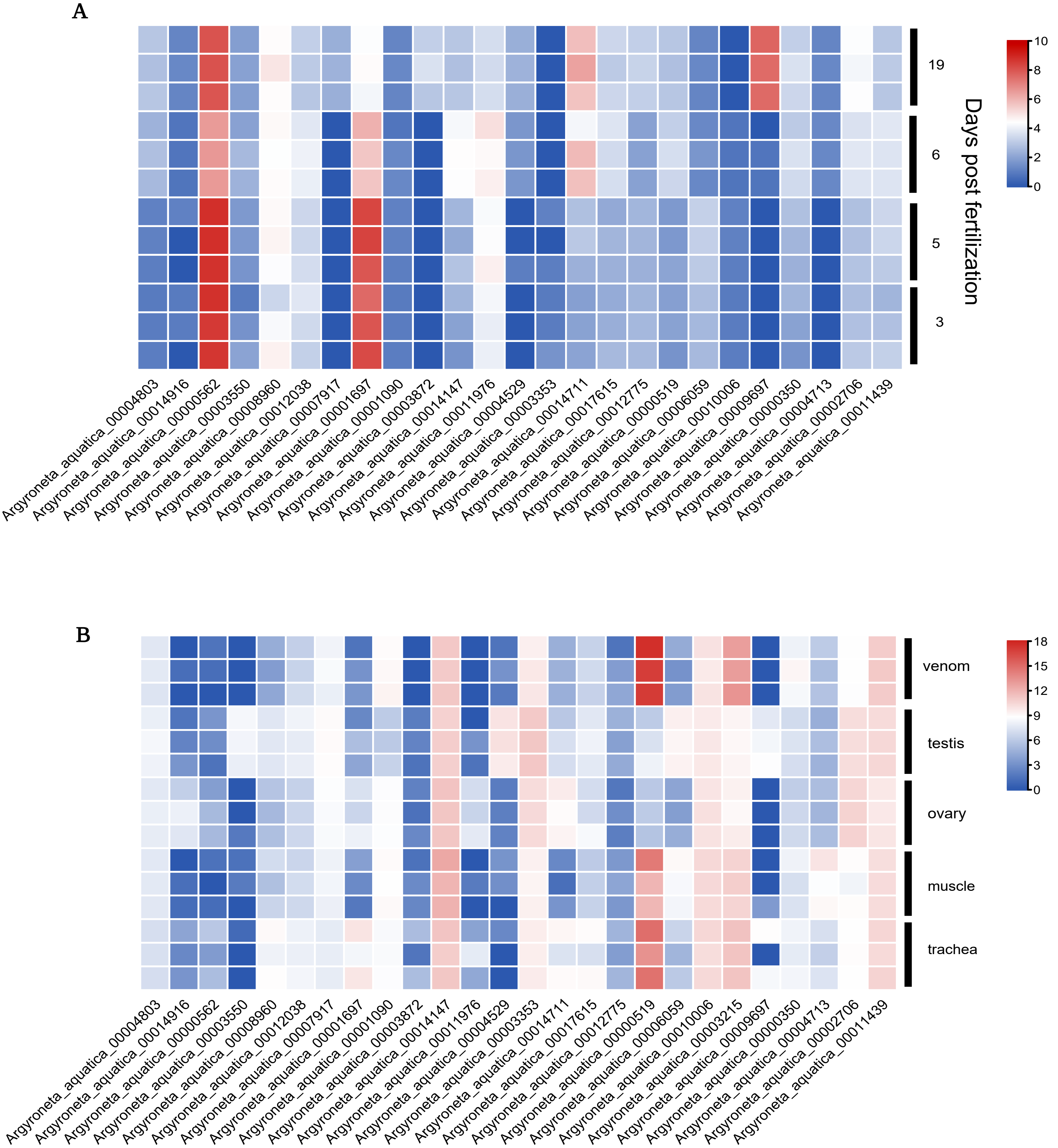


**Figure S16. The expression of aquatic positive selection genes among different tissues and develop stages.**
A. The expression of aquatic positive genes among 3, 5, 6 and 19 days post fertilization.
B. The heatmaps of aquatic positive genes expression among different tissues including venom, trachea, muscle, testis and ovary.

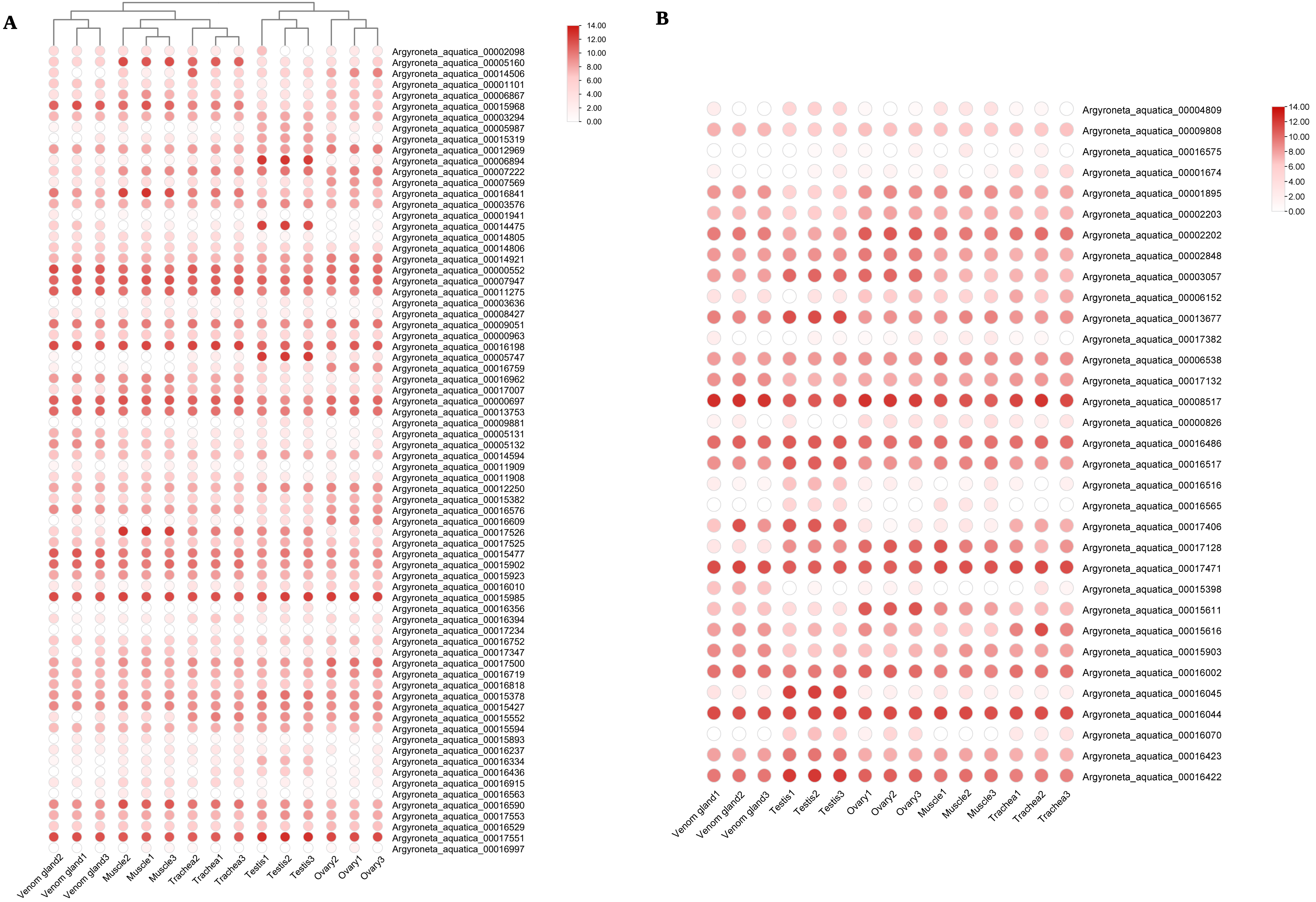


Figure S17. The expression of ABC (A) and ACAD (B) genes in different tissues of water spider.

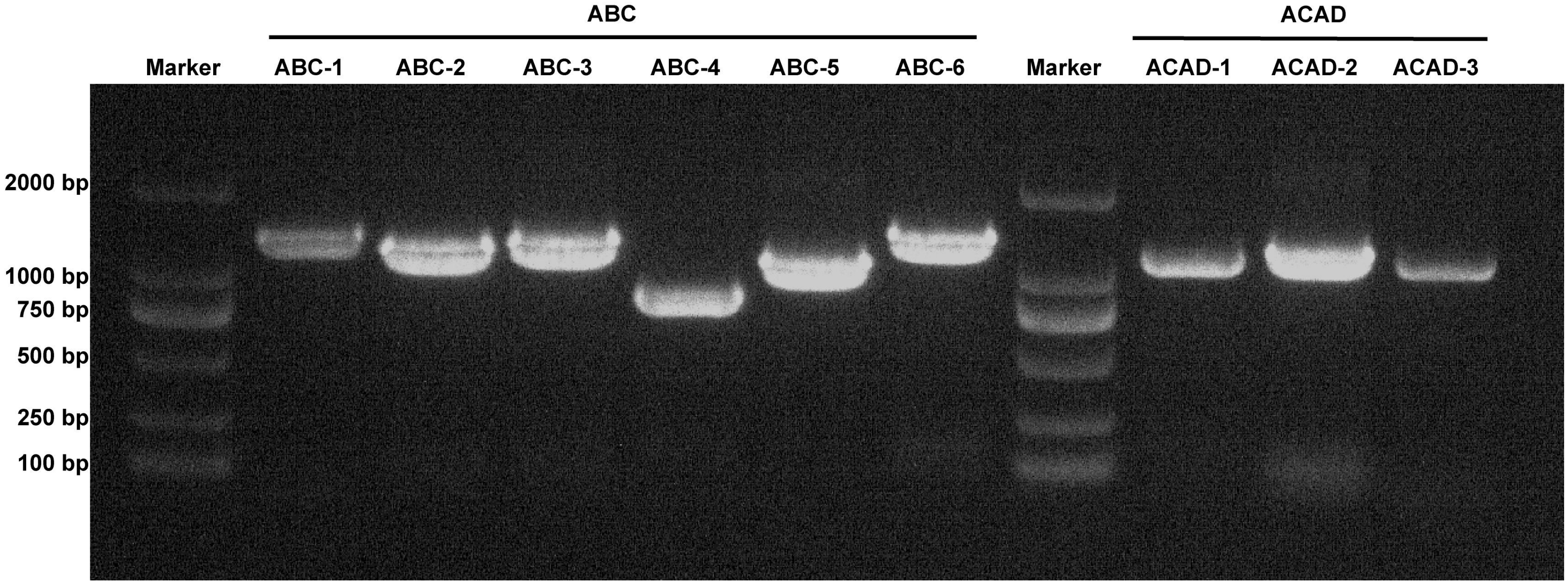


Figure S18. PCR identified of nine HGT genes in water spider genome, including six ABC and three ACAD genes.

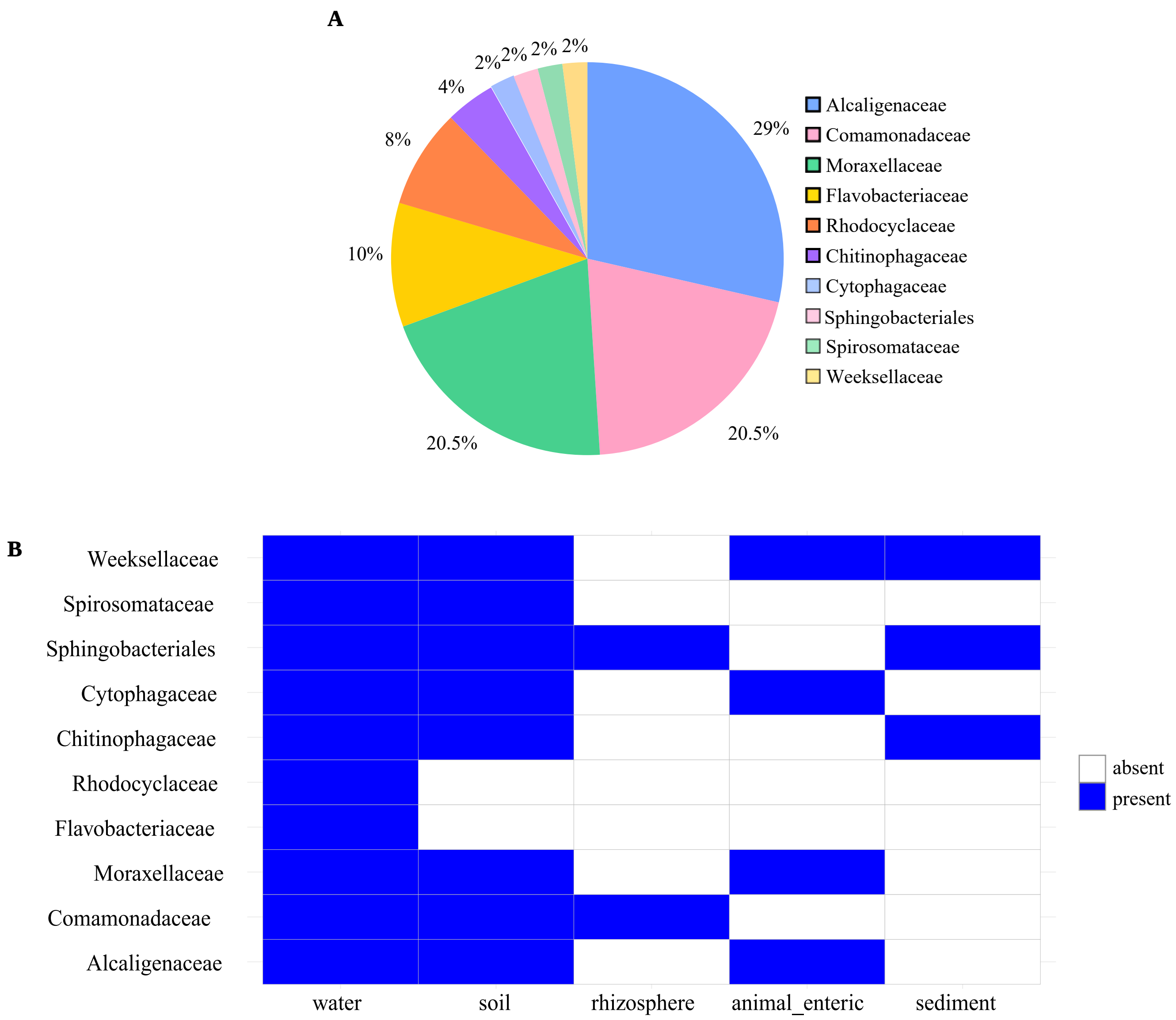


**Figure S19. The source of HGT-acquired ABC and ACAD genes in water spider.**
A. The genus of bacteria of the source for HGT-acquired ABC and ACAD genes.
B. The main living environment which transferred for water spider HGT-acquired ABC and ACAD genes.

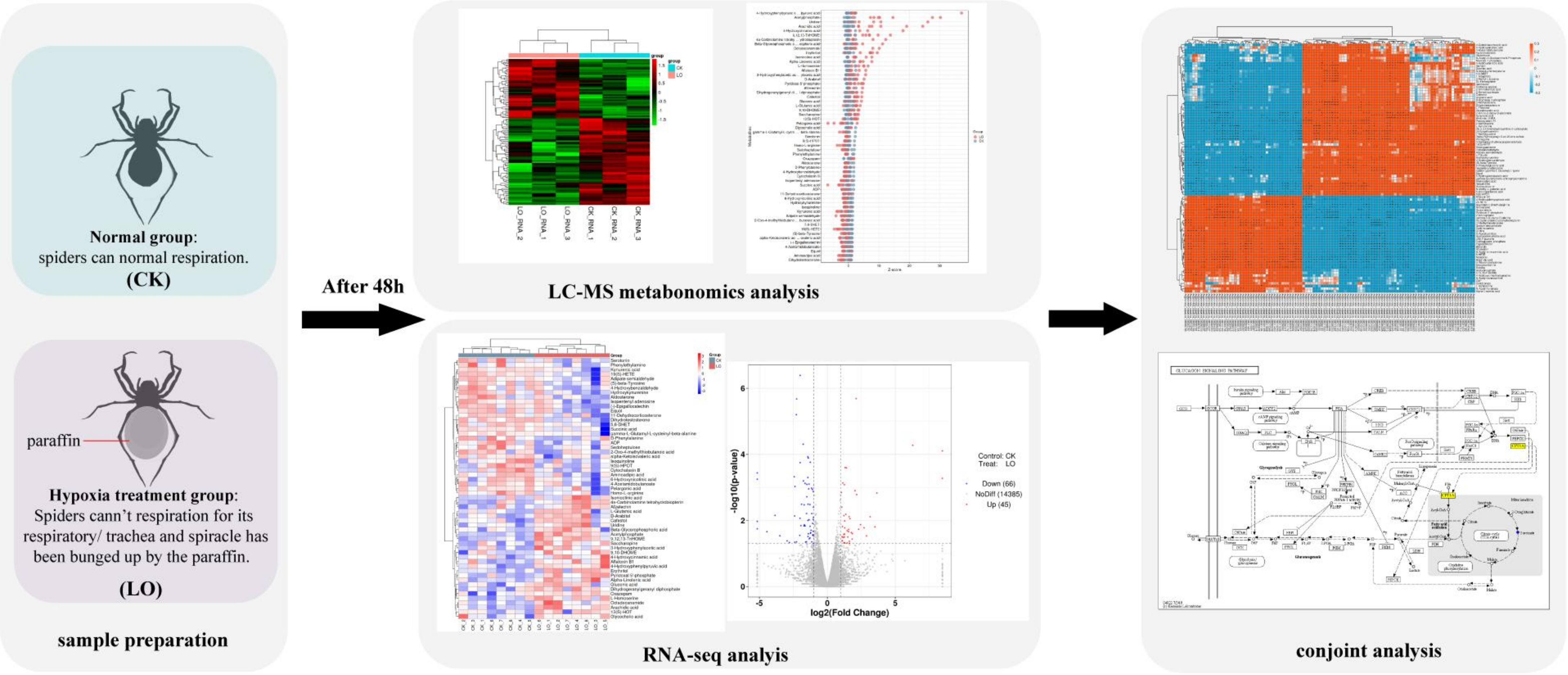


Figure S20. The process of metabonomics and tanscriptomics analysis with water spider hypoxia treatment.
